# A tissue-specific collaborative mixed model for jointly analyzing multiple tissues in transcriptome-wide association studies

**DOI:** 10.1101/789396

**Authors:** Xingjie Shi, Xiaoran Chai, Yi Yang, Qing Cheng, Yuling Jiao, Jian Huang, Can Yang, Jin Liu

## Abstract

Transcriptome-wide association studies (TWAS) integrate expression quantitative trait loci (eQTLs) studies with genome-wide association studies (GWASs) to prioritize candidate target genes for complex traits. Several statistical methods have been recently proposed to improve the performance of TWAS in gene prioritization by integrating the expression regulatory information imputed from multiple tissues, and made significant achievements in improving the ability to detect gene-trait associations. The major limitation of these methods is that they cannot be used to elucidate the specific functional effects of candidate genes across different tissues. Here, we propose a tissue-specific collaborative mixed model (TisCoMM) for TWAS, leveraging the co-regulation of genetic variations across different tissues explicitly via a unified probabilistic model. TisCoMM not only performs hypothesis testing to prioritize gene-trait associations, but also detects the tissue-specific role of candidate target genes in complex traits. To make use of widely available GWAS summary statistics, we extend TisCoMM to use summary-level data, namely, TisCoMM-S^2^. Using extensive simulation studies, we show that type I error is controlled at the nominal level, the statistical power of identifying associated genes is greatly improved, and false positive rate (FPR) for non-causal tissues is well controlled at decent levels. We further illustrate the benefits of our methods in applications to summary-level GWAS data of 33 complex traits. Notably, apart from better identifying potential trait-associated genes, we can elucidate the tissue-specific role of candidate target genes. The follow-up pathway analysis from tissue-specific genes for asthma shows that the immune system plays an essential function for asthma development in both thyroid and lung tissues.

## Introduction

Over the last decade, GWASs have achieved remarkable successes in identifying genetic susceptible variants for a variety of complex traits [1]. However, the biological mechanisms to understand these discoveries remain largely elusive as majority of these discoveries are located in non-coding regions [2]. Recent expression quantitative trait loci (eQTLs) studies indicate that the expression regulatory information may play a pivotal role bridging both genetic variants and traits [3, 4, 5]. Cellular traits in comprehensive eQTL studies can serve as reference data, providing investigators with an opportunity to examine the regulatory role of genetic variants on gene expression. For example, the Genotype-Tissue Expression (GTEx) Project [6] has provided DNA sequencing data from 948 individuals and collected gene-expression measurements of 54 tissues from these individuals in the recent V8 release.

Transcriptome-wide association studies (TWAS) has been widely used to integrate the expression regulatory information from these eQTL studies with GWAS to prioritize genome-wide trait-associated genes [7, 8, 9]. A variety of TWAS methods have been proposed using different prediction models for expression imputation, including the parametric imputation models, e.g., PrediXcan [7], TWAS [8], CoMM [10] and CoMM-S^2^ [11], and the nonparametric imputation model, e.g., Tigar [12]. These methods have been used for analyzing many complex traits with expression profiles from different tissues, successfully enhancing the discovery of genetic risk loci for complex traits [13, 9]. To further improve the power of identifying potential target genes, two recent studies were proposed by leveraging the substantial shared eQTLs across different tissues, i.e., MultiXcan [14] and UTMOST [15]. They use a step-wise procedure by first conducting imputation for gene expressions across multiple tissues and then performing subsequent association analysis using a multivariate regression that pools information across different tissues. Compared to single-tissue methods, these multi-tissue strategies enhance the imputation accuracy for gene expression and thus improve the power of identifying potential target genes.

Despite their successes, the existing multi-tissue methods have several limitations. First, MultiXcan and UTMOST cannot be used to identify the tissue-specific gene-trait associations. Many studies have shown that genes associated with complex traits are always regulated in a tissue-specific manner [16, 17, 18, 9]. For example, a recent study across 44 tissues confirmed this phenomenon in 18 complex traits [19], implying the persuasive role of tissue-specific regulatory effects in a wide range of complex traits. Using a single-tissue test, one can easily reach a false conclusion regarding which tissue that a gene affects traits through. Second, both MultiXcan and UTMOST rely on a step-wise inference framework, ignoring the uncertainty in the process of expression imputation and thus losing power, especially when cellular-heritability is small [10]. Recently, CoMM [10] and its variant for summary-level data, CoMM-S^2^ [11], have been proposed to account for uncertainty in the process of expression imputation. Third, MultiXcan and UTMOST do not make efficient use of the shared patterns of eQTLs across tissues, where MultiXcan uses principal component analysis (PCA) regularization on the predicted expression data, and UTMOST uses penalized regularization on coefficients for eQTL effects. A study of GTEx revealed these shared patterns [20], and later many efforts have been made to take advantage of them in the analysis for GTEx data. For example, Urbut et al. proposed statistical methods for estimating and testing eQTL effects explicitly incorporating this extensively tissue-shared patterns [21], shedding light on how to account for the tissue-shared eQTLs in statistical modeling successfully.

To overcome these limitations, we propose a tissue-specific collaborative mixed model (TisCoMM) for TWAS, providing a principled way to perform gene-trait joint and tissue-specific association tests across different tissues. Our method allows us not only to perform hypothesis testing to prioritize gene-trait association but also to uncover the tissue-specific role of candidate genes. By conditioning on the trait-relevant tissues, one could largely remove the spurious associations due to highly correlated gene expressions among multiple tissues. As a unified model, TisCoMM jointly conducts the “imputation” and the association analysis, pooling expression regulatory information across multiple tissues explicitly. Furthermore, we extend TisCoMM to use summary statistics from a GWAS, namely, TisCoMM-S^2^. In simulations, we show that both TisCoMM and TisCoMM-S^2^ provide correctly controlled type I error and are more powerful than existing multi-tissue methods. More importantly, our methods can be used to test for the tissue-specific role of candidate genes. We illustrate the benefits of our methods using summary-level GWAS data in 33 complex traits. Results show that our findings have biologically meaningful implications. The follow-up pathway analysis from tissue-specific genes for asthma shows that the regulated immune system in both thyroid and lung tissues could have significant impact on asthma development.

## Results

### Method overview

Our method, TisCoMM, jointly integrates expression regulatory information across multiple tissues by considering two models. The first one models the relationship between genetic factors and gene expressions across multiple tissues in the eQTL data set,

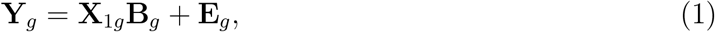

where 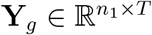 is expression matrix of *n*_1_ samples across *T* tissues for gene *g*, 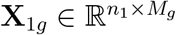 is the standardized genotype matrix corresponding to *M*_*g*_ nearby single nucleotide polymorphisms (SNPs) of gene *g* in the eQTL data, **B**_*g*_ is an *M*_*g*_ × *T* matrix of the corresponding effect sizes across *T* tissues and **E**_*g*_ is an *n*_1_ × *T* matrix for random errors from a multivariate normal distribution 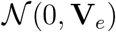. Here, **V**_*e*_ captures the correlations among tissues from the same individual. Then we assume that phenotypic value **z** and standardized genotype **X**_2*g*_ in GWAS are related by

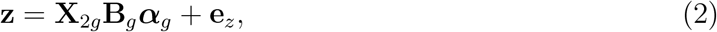

where **z** is an *n*_2_ × 1 vector of phenotypic values, 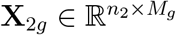 is the standardized genotype matrix corresponding to *M*_*g*_ nearby variants of gene *g* in the GWAS data, ***α***_*g*_ is a *T* × 1 unknown parameter vector of interest that represents the effect sizes of “imputed” gene expression across *T* tissues for gene *g*, and 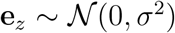 is an *n*_2_ × 1 vector of independent errors associated with the trait. Our TisCoMM can be depicted as Figure 1, within which Figure 1A illustrates the TisCoMM method combing both the expression prediction model (1) and the corresponding association model (2) together with data input and output.

**Figure 1:**
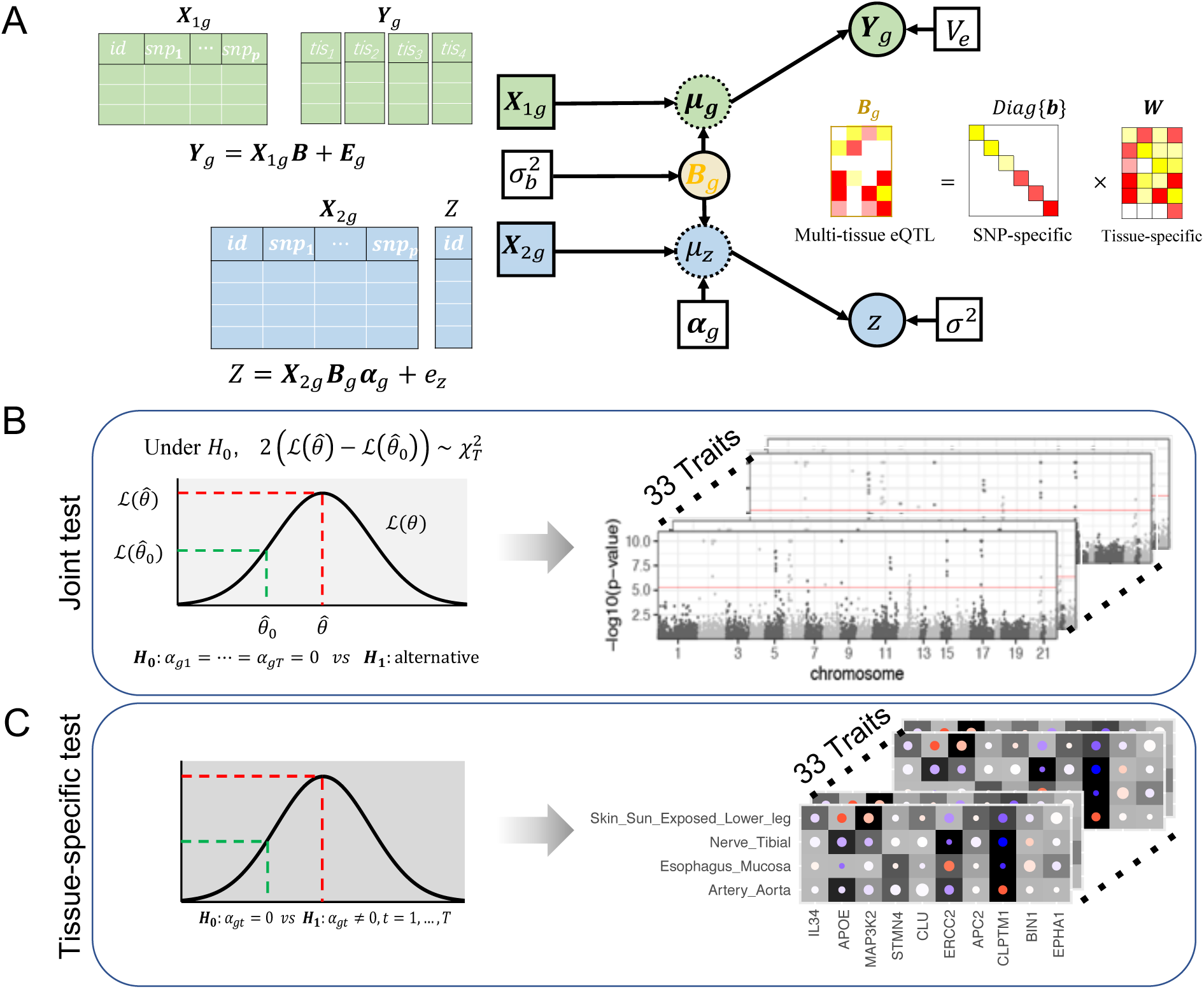
TisCoMM workflow. **A.** Two sets of TisCoMM input matrices are highlighted in green and blue separately (left). The probabilistic graphical model for TisCoMM is shown in the middle, which integrates gene expressions and models the co-regulation of cis-SNPs across different tissues explicitly. *μg* and *μ*_*z*_ denote expectations of gene expression in eQTL and phenotype in GWAS, respectively. The decomposition of the **B** matrix is illustrated on the right-hand side of the figure. **B.** The TisCoMM joint test for all genes to prioritize candidate causal genes. See more details of 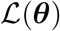 in Methods section. The example outputs (right) are shown as Manhattan plots for 33 traits. **C.** The TisCoMM tissue-specific test for all candidate genes to explore the tissue-specific roles of candidate genes. The example outputs (right) are shown as heatmaps which summarize the tissue-specific effect of each gene. Significance level, effect size, and heritability are converted into background color, circle color, and circle size.

To pool expression regulatory information across relevant tissues, we assume the factorizable assumption [22, 23] for **B**_*g*_ = [*β*_*jt*_], *j* = 1, …, *M*_*g*_, *t* = 1, …, *T*. This assumption has been empirically validated for GTEx data in an imputation study [24] and Park et al. further used this assumption in a multi-tissue TWAS [25]. Here, we assume that the effect size of cis-SNP *j* in tissue *t* can be factorized by variant-dependent and tissue-dependent components: *β*_*jt*_ = *b*_*j*_*w*_*jt*_, where *b*_*j*_ (variant) is the eQTL effect of cis-SNP *j* shared in all the *T* tissues, and *w*_*jt*_ is the tissue-specific effect size. Thus, we have **B**_*g*_ = diag{**b**}**W**. This factorization allows us to model the co-regulation of cis-SNPs shared across different tissues explicitly (Figure 1A, right). To make TisCoMM identifiable, we further assume that *b*_*j*_ independently follows a normal distribution 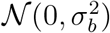 due to polygenicity and by following the adaptive weighting strategy used in [24], the adaptive weight *w*_*jt*_ is estimated using the marginal regression of gene expression in tissue *t* on the *j*-th genetic variant.

The parameter of our interest in TisCoMM is the vector of effect size ***α***_*g*_. To prioritize candidate target genes, we conduct hypothesis testing for a joint null, *H*_0_ : ***α***_*g*_ = 0 (Figure 1B). To further explore the tissue-specific roles of candidate genes, we conduct hypothesis testing for each tissue, *H*_0_ : *α*_*gt*_ = 0, *t* = 1, …, *T* (Figure 1C). We refer to the two inference tasks as the TisCoMM joint test and TisCoMM tissue-specific test, respectively. We develop an expectation-maximization (EM) algorithm for parameter estimation by maximizing the complete-data likelihood. A parameter expansion technique is further adopted to accelerate computational efficiency (see details in Supplementary Text). In contrast to the existing two-step TWAS methods, we perform TisCoMM analysis in a unified model by treating **b** as a hidden random variable. Generally, the computational cost for the TisCoMM tissue-specific test is 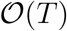 of that for the TisCoMM joint test. To enable computational efficiency, we only conduct the TisCoMM tissue-specific test for candidate genes detected in the joint test, rather than for all genes.

In a single-tissue analysis, it is difficult to explore the tissue-specific role of a candidate gene. The disease-associated genes will be identified in all the causal tissues as well as the tissues (possibly non-causal) highly correlated with the causal one, because there exist sharing patterns for expressions in multiple tissues. By conditioning on the trait-relevant tissues, our tissue-specific test could largely remove the spurious discoveries due to correlated expression across tissues.

### Inferring TisCoMM results from GWAS summary statistics

To make our method widely applicable, we extend TisCoMM to use summary-level GWAS data, denoted as TisCoMM-S^2^. The model details are given in Supplementary Text.

We observe high concordance between TisCoMM and TisCoMM-S^2^ results. Figure 2 shows the comparison of TisCoMM and TisCoMM-S^2^ test statistics for ten traits from the Northern Finland Birth Cohorts program 1966 (NFBC1966) data set [26] (see Methods section). The reference panel was 400 subsamples from the NFBC1966 data set. The high correlation between TisCoMM and TisCoMM-S^2^ suggests the goodness of detections for trait-associated genes using summary-level GWAS data.

**Figure 2:**
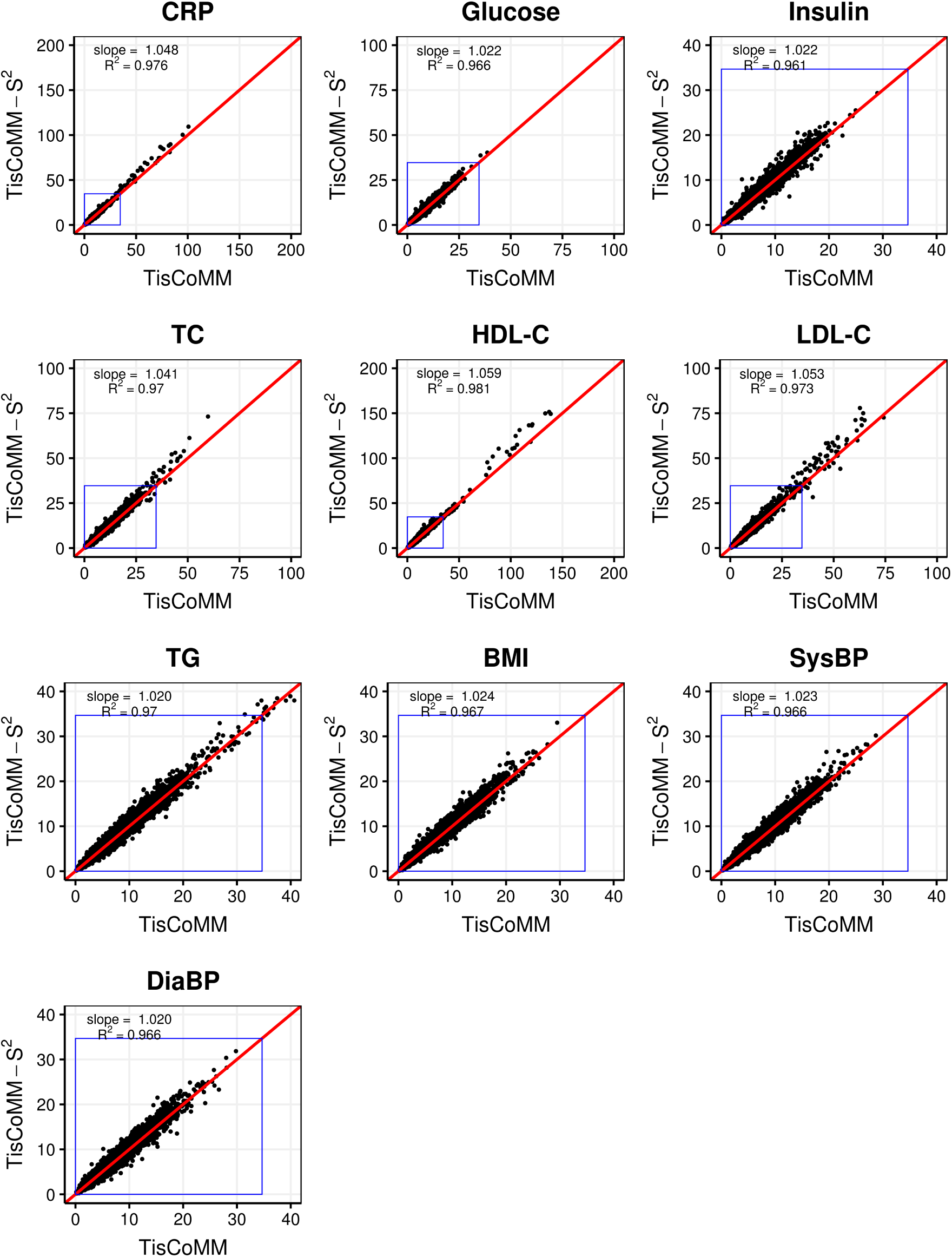
Comparison of TisCoMM and TisCoMM-S^2^ results in NFBC1966 traits. The reference panel is subsamples from the NFBC1966 data set. The summary-based method shows similar results to the individual-based method. The blue rectangle indicates the null region.

To test the robustness of TisCoMM-S^2^, we applied European subsamples from 1000 Genomes as the reference panel. Note that the NFBC1966 data set is Finns study, and it is well known that Finns have significant genetic differences with other Europeans [27]. Hence, the estimated LD did not well match that of the GWAS study. Supplementary Figure S1 shows the performance of TisCoMM-S^2^ using European subsamples as a reference panel data set. Despite the high concordance between TisCoMM and TisCoMM-S^2^ in the null region (Λ > 34.67 = *p*-values > 5 × 10^−6^), the test statistics of TisCoMM-S^2^ in the non-null region are much more significant than TisCoMM.

### Simulation

#### Methods for comparison

To detect gene-trait association, we compared the performance of three methods in the main text: (1) our TisCoMM and TisCoMM-S^2^ implemented in the R package *TisCoMM*; (2) MultiXcan and S-MultiXcan implemented in the MetaXcan package available at http://gene2pheno.org/; (3) UTMOST available at https://github.com/Joker-Jerome/UTMOST/. To detect the tissue-specific effect, we compared the performance of Tis-CoMM tissue-specific test with three single-tissue methods that include (1) CoMM available at https://github.com/gordonliu810822/CoMM; (2) PrediXcan available at http://gene2pheno.org/; (3) TWAS relies on the BSLMM [28] implemented in the GEMMA [28] software. All methods were used with default settings. We conducted comprehensive simulations to gauge the performance of each method better by performing gene-trait joint and tissue-specific tests across different tissues.

#### Simulation settings

In detail, we considered the following simulation settings. We set {*n*_1_, *n*_*r*_, *n*_2_} = {400; 400; 5, 000} as the sample size for eQTL data, GWAS data and reference panel data. We first generated the genotype data for *M*_*g*_ = 400 cis-SNPs from a multivariate normal distribution assuming an autoregressive correlation with parameter *ρ*. We then discretized each SNP to a trinary variable {0, 1, 2} by assuming Hardy-Weinberg equilibrium and a minor allele frequency randomly selected from a uniform [0.05, 0.5] distribution. The genotype correlation was varied at *ρ* = {0.2, 0.5, 0.8}. All three genotype matrices, **X**_1*g*_, **X**_*rg*_, and **X**_2*g*_, for eQTL data, GWAS data and reference panel data, respectively, are generated in this manner.

To generate multi-tissue gene expressions, we considered different cellular-level heritability levels 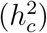 and sparsity levels (*s*). These are key parameters to describe the genetic architecture of gene expression [29]. The cellular-level heritability represents the proportion of variance of the gene expression that can be explained by genotype, while sparsity represents the proportion of genetic variants that are associated with the gene expression. First, SNP effect size **B**_*g*_ = diag{**b**}**W** is generated. Specifically, we simulated SNP effect size **b** from a standard normal distribution, and randomly selected 10%, 50% or 100% of the SNPs to have non-zero tissue-specific effect **W** for gene expressions in all *T* tissues, while simulated their effects from a standard normal distribution. We then simulated errors **E**_*g*_ from a normal distribution, where their variances were chosen according to 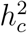, and the covariance structure was autoregressive with *ρ*_*e*_ = 0.5. Here we set 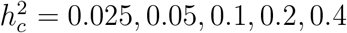. Afterward, we simulated a multi-tissue eQTL data set assuming **Y**_*g*_ = **X**_1*g*_**B**_*g*_ + **E**_*g*_.

To simulate a quantitative trait, we generated nonzero entries of ***α***_*g*_ from a uniform distribution and **e**_*z*_ from a normal distribution. The variance *σ*^2^ was chosen according to the tissue-level heritability 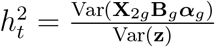. Here we set 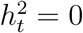 for null simulations and type I error control examination and 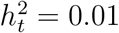 for non-null simulations and power comparisons.

#### Simulation I: Testing gene-trait associations

We focus on the detection of trait-associated genes in the first set of simulations. Here, we compared TisCoMM and TisCoMM-S^2^ with three different multi-tissue methods that include MultiXcan, S-MultiXcan, and UTMOST. We set *T* = 10, and all tissues are causal. For each scenario, we run 5,000 replicates. We first examined type I error control of different methods under the null. Results are shown in Supplementary Figures S2 – S6. By comparing the distribution of *p*-values with the expected uniform distribution, we observe that all methods provide well-controlled type I errors.

Next, we examined the power of different methods under the alternative hypothesis, as shown in Figure 3. We observe that the performance of all five methods improves with the increment of cellular heritability. In general, the summary-level methods (TisCoMM-S^2^ and S-MultiXcan) perform similarly to their counterparts in individual-level data. Moreover, TisCoMM and TisCoMM-S^2^ have better performance than other alternative methods when cellular heritability is relatively small 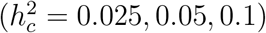, and comparable performance when cellular heritability is large. Finally, we observe that although our model favors dense eQTLs, it was robust to the sparsity level *s*. Specifically, the power of TisCoMM and TisCoMM-S^2^ in the setting where 10% of cis-SNPs have non-zero effects on gene expression are similar to the setting where all cis-SNPs have non-zero effects.

**Figure 3:**
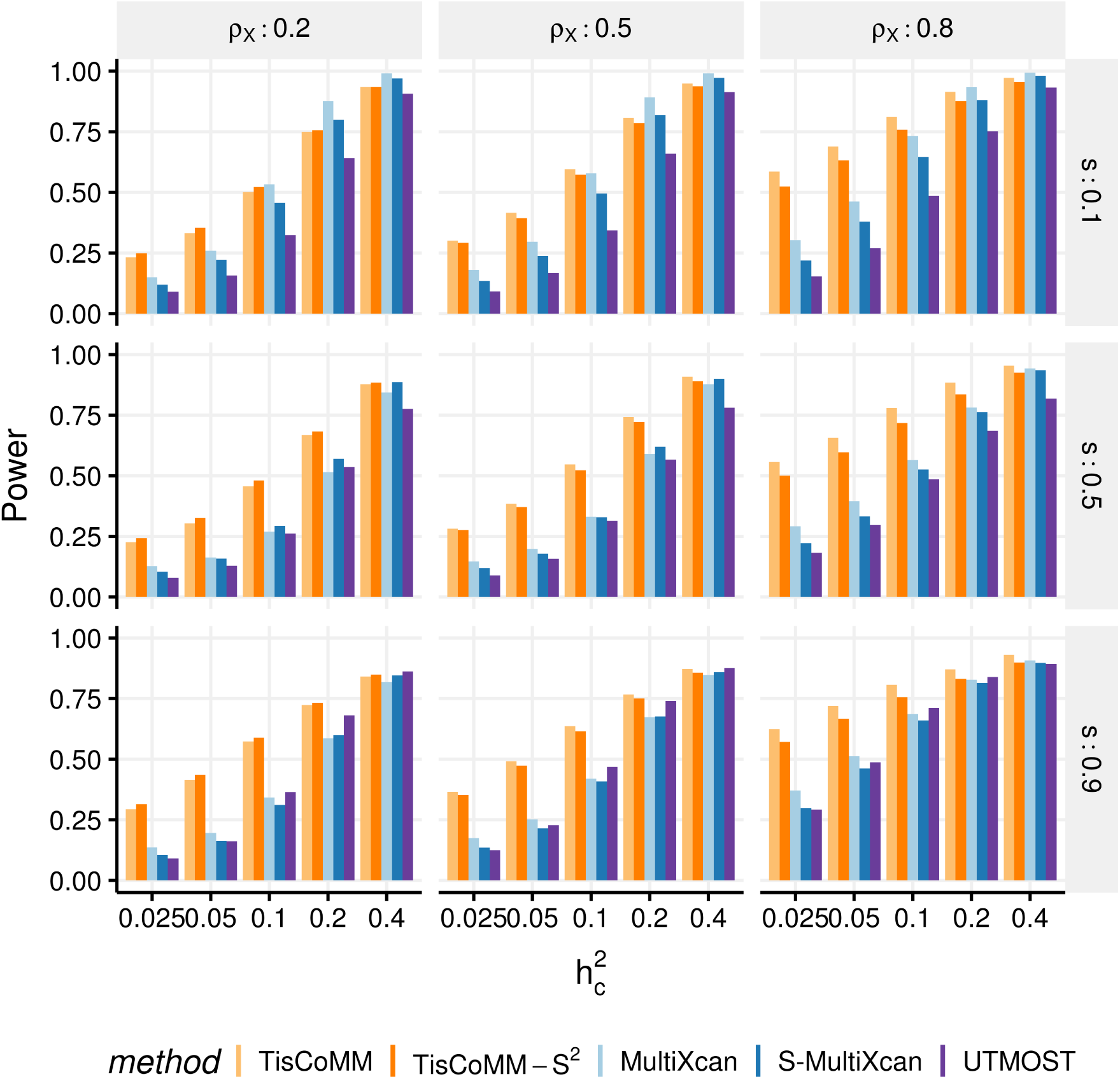
TisCoMM joint test outperforms the other multi-tissue methods. The number of replicates is 5,000. In each subplot, the x-axis stands for the SNP heritability level, and the y-axis stands for the proportion of significant genes within 5,000 replicates.

#### Simulation II: Testing tissue-specific effects

We focus on the detection of tissue-specific effects in the second set of simulations. Here, we compared the TisCoMM tissue-specific test with the single-tissue methods including CoMM [10], PrediXcan [7], and TWAS[8] under the alternative hypothesis with fixed tissue heritability 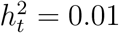 and fixed sparsity *s* = 0.1. We considered three tissues *T* = 3 and varied the number of causal tissues to simulate different levels of tissue specificity of a trait. Specifically, we considered settings with one (*α*_*g*2_ = *α*_*g*3_ = 0) and two causal tissues (*α*_*g*3_ = 0), respectively. To allow correlated gene expression in the GWAS, the nonzero of tissue-specific effect **W** was generated with rows drawn from a multivariate normal distribution, with AR correlation parameter *ρ*_*W*_ = 0.2, 0.5, 0.8. A large value of *ρ*_*W*_ implies a higher correlation among columns of **X**_2*g*_**B**_*g*_. Other sittings are similar to Simulation I.

We repeated the whole process 1,000 times. We calculated statistical power and false positive rate (FPR) as the proportion of *p*-values reaching the significance level in causal tissues and non-causal tissues, respectively. Specifically, we set the significance level at 0.05/3 for all considered methods. Figure 4 shows simulation results for the case that one tissue is causal. We observe that in all settings, the TisCoMM tissue-specific test has comparable or slightly inferior power, as shown in Figure 4A, compared to the single-tissue methods, but much smaller FPR (Figure 4B). As expected, the statistical power of all methods increases with cellular heritability 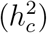. However, the FPR of single-tissue methods substantially inflates while that of TisCoMM tissue-specific test remains at the same level. Furthermore, the FPR of TisCoMM tissue-specific test does not vary with correlations among expressions across multiple tissues (*ρ*_*W*_) while that of single-tissue methods increase with *ρ*_*W*_. The similar pattern could be observed for the case that two tissues are causal (Supplementary Figure S7). These results demonstrate the usefulness of TisCoMM tissue specific test in exploring the tissue-specific role of genes.

**Figure 4:**
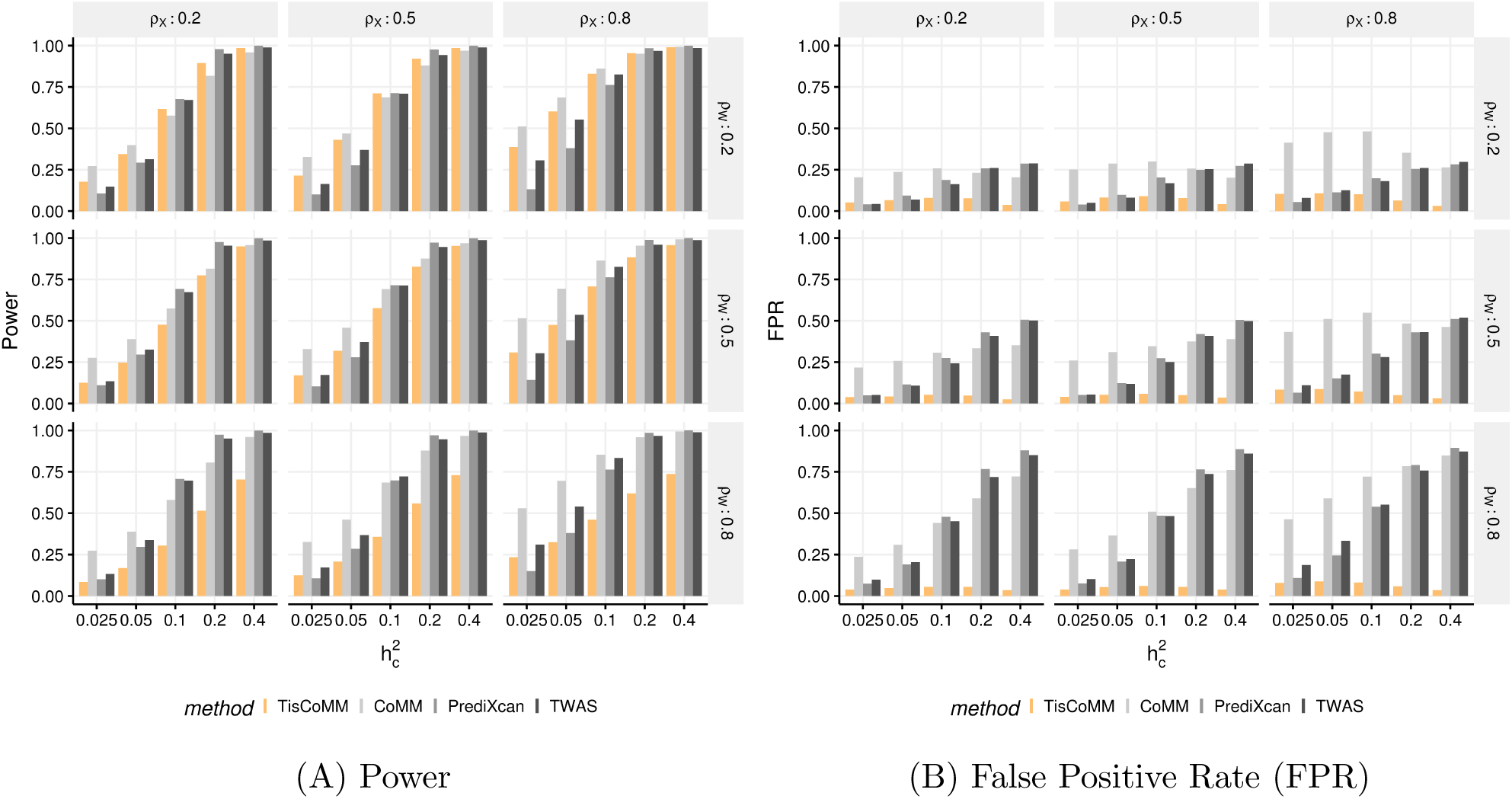
The comparison of the TisCoMM tissue-specific test and the single-tissue association tests under the alternative hypothesis with one causal tissue. **A.** The power of TisCoMM tissue-specific test and the single tissue methods with Bonferroni correction applied. **B.** The corresponding false positive rates under each setting.

### Real Data Applications

We performed multi-tissue TWAS analysis for summary-level GWAS data in 33 complex traits (see Supplementary Table S1 for details), including 15 traits from Gamazon et al. [19] and 18 traits from the UK Biobank. Hereafter we refer to as NG traits and UKB traits, respectively. These traits can be roughly divided into four categories, including metabolites (e.g., HDL-C, LDL-C and fasting glucose), autoimmune diseases (e.g., asthma, Crohn’s disease and macular degeneration), psychiatric/neurodegenerative disorders (e.g., Alzheimer’s disease, major depression disorder, and psychiatric disorder), and cardiovascular disorders (e.g., coronary artery disease and peripheral vascular disease). The Genotype-Tissue Expression (GTEx) Project [6] reported eQTL in 48 tissues, where the number of genes in each tissue ranges from 16,333 to 27,378. In the analysis, we extracted cis-SNP that are within either 500 kb upstream of the transcription start site or 500 kb downstream of the transcription end site.

In a single-tissue analysis, there are two different strategies to select a tissue for TWAS: one uses expressions from the most biologically related tissue while the other selects a tissue with the largest number of available individuals [9]. To select multiple tissues for TisCoMM-S^2^, there exists a trade-off between biological relevance and its corresponding sample size for each tissue. In [19], it provides the most biologically related tissues and thus we used trait-relevant tissues for the NG traits from Supplementary Table 2 in [19]. In detail, for each trait, a set of tissues with significant enrichment *p*-values (after Bonferroni correction) was identified, and a subset with more than 100 overlapped samples [30] was chosen for further analysis in TisCoMM-S^2^. On the other hand, although methods like LD score regression [17] can be used for the UKB traits, it is difficult to balance the tissue relevance and sample size for each tissue. To make efficient use of the GTEx data set, we used six tissues with the largest number of overlapped samples for the UKB traits.

The analysis for each trait based on its GWAS summary statistics together with the eQTL data from multiple tissues can be done around 100 min on a Linux platform with 2.6 GHz Intel Xeon CPU E5-2690 with 30720 KB cache and 96 GB RAM (0nly 10~12 GB RAM used) on 24 cores.

### TisCoMM-S^2^ joint test provides statistically powerful results of disease relevant genes

To prioritize trait-associated genes, we compared TisCoMM-S^2^ with other two multi-tissue TWAS methods, i.e., S-MultiXcan and UTMOST. Both alternative methods take advantage of prediction models to impute gene expressions. The prediction models used here were Elastic Net models trained on 48 GTEx tissues. See Table 1 and 2 for the summary of detections across different approaches for the 15 NG and 18 UKB traits, respectively. Generally, TisCoMM-S^2^ identifies more genome-wide associations than S-MultiXcan and UTMOST in most traits. In detail, TisCoMM-S^2^/S-MultiXcan/UTMOST identified 3,058/2,008/1,769, and 443/338/277 genome-wide significant genes in all the NG traits and UKB traits, respectively. Their qq-plots of *p*-values are shown in Supplementary Figures S8 – S11 and plots for their genomic inflation factors are shown in Supplementary Figure S12. As case study examples, we carefully examined the results for late-onset Alzheimer’s disease (LOAD) and asthma.

**Table 1:**
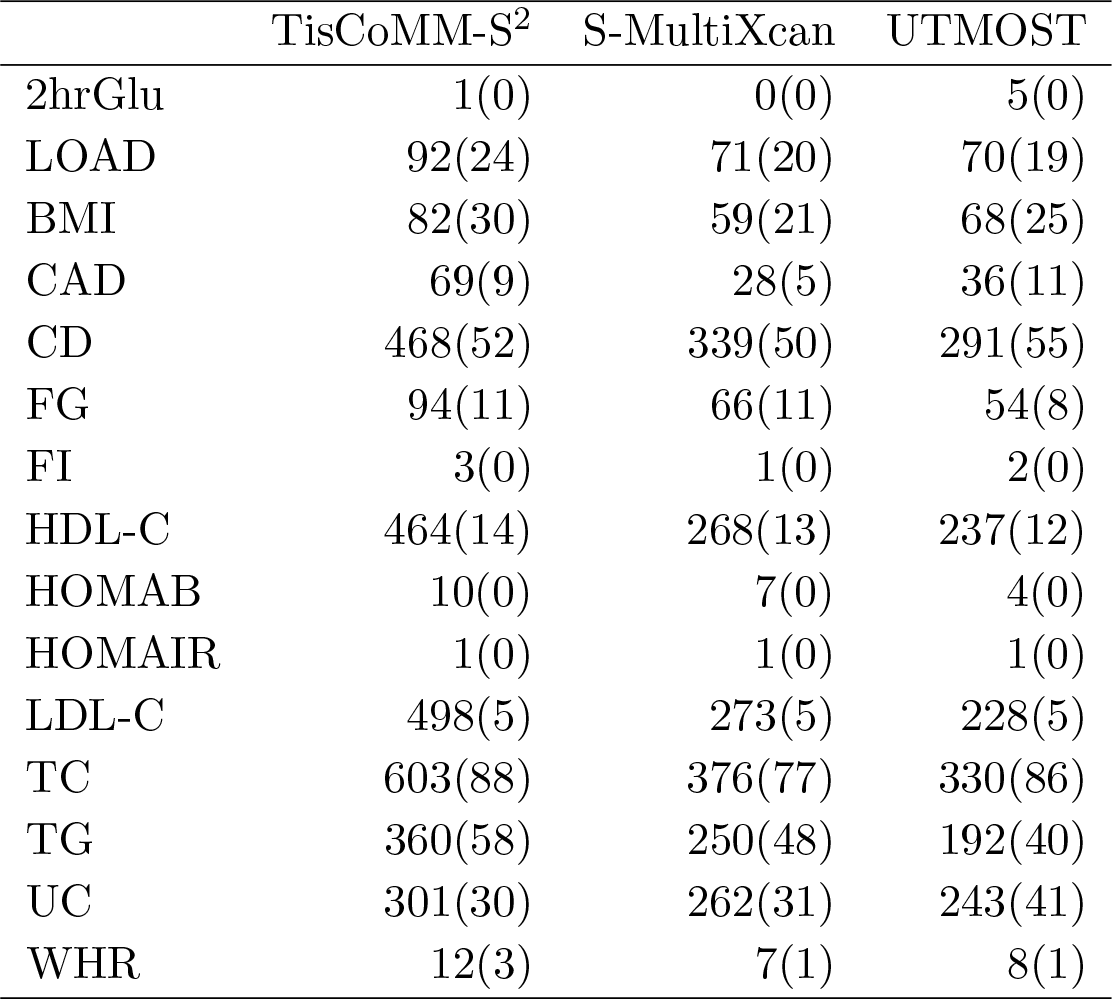
Numbers of significant gene-trait associations across 15 NG traits. The reference penal is European subsamples from 1000 Genome. The number in the parenthesis denoted genes reported on the GWAS catalog. The full names of traits can be found in Supplementary Table S1.

|  | TisCoMM-S<sup>2</sup> |  | S-MultiXcan | UTMOST |
| --- | --- | --- | --- | --- |
| 2hrGlu | 1(0) |  | 0(0) | 5(0) |
| LOAD | 92(24) |  | 71(20) | 70(19) |
| BMI | 82(30) |  | 59(21) | 68(25) |
| CAD | 69(9) |  | 28(5) | 36(11) |
| CD | 468(52) |  | 339(50) | 291(55) |
| FG | 94(11) |  | 66(11) | 54(8) |
| FI | 3(0) |  | 1(0) | 2(0) |
| HDL-C | 464(14) |  | 268(13) | 237(12) |
| HOMAB | 10(0) |  | 7(0) | 4(0) |
| HOMAIR | 1(0) |  | 1(0) | 1(0) |
| LDL-C | 498(5) |  | 273(5) | 228(5) |
| TC | 603(88) |  | 376(77) | 330(86) |
| TG | 360(58) |  | 250(48) | 192(40) |
| UC | 301(30) |  | 262(31) | 243(41) |
| WHR | 12(3) |  | 7(1) | 8(1) |

**Table 2:**
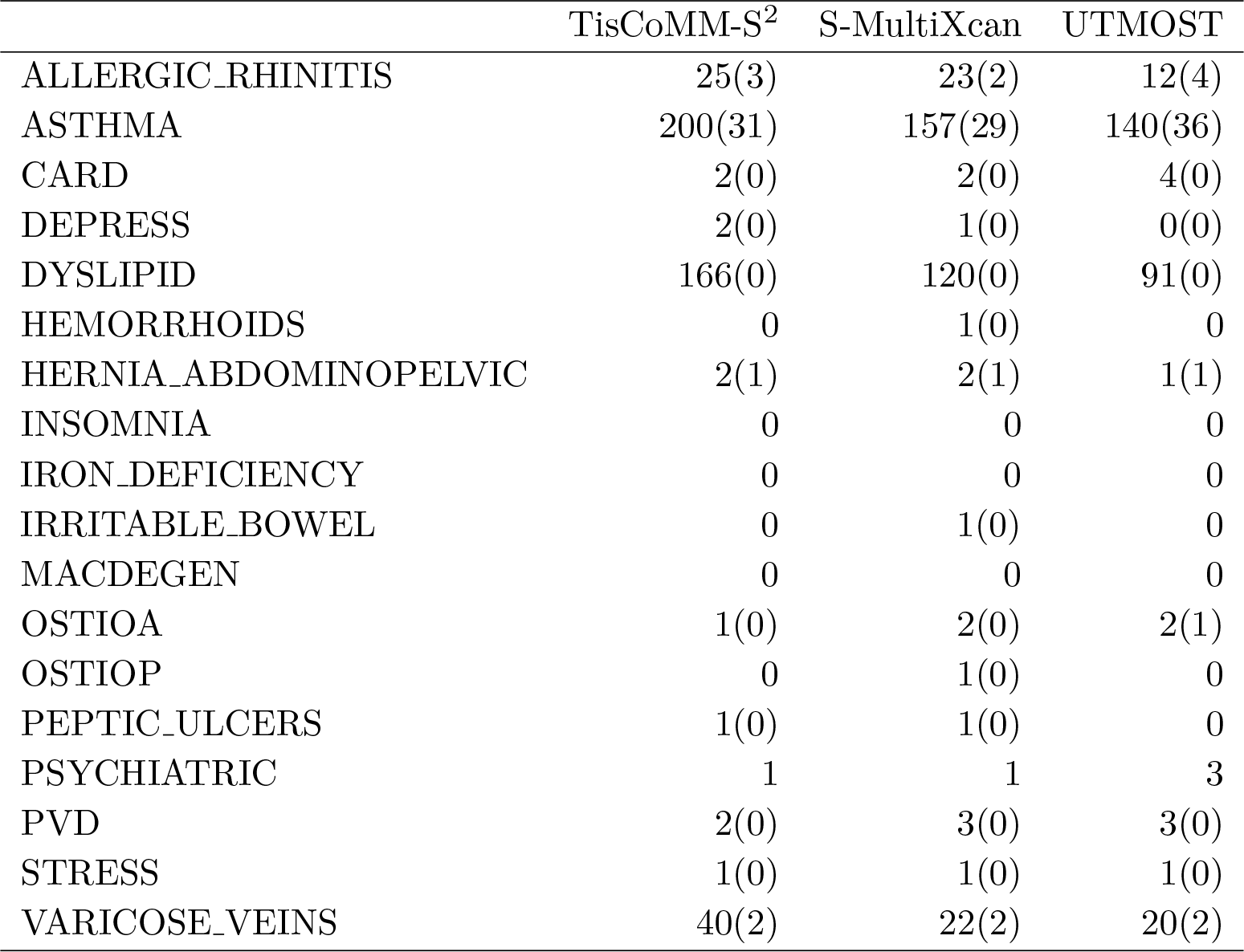
Numbers of significant gene-trait associations across 18 UKB traits. The reference penal data is European subsamples from 1000 Genome. The number in the parenthesis denoted genes reported on the GWAS catalog. The full names of traits can be found in Supplementary Table S1.

|  | TisCoMM-S<sup>2</sup> |  | S-MultiXcan | UTMOST |
| --- | --- | --- | --- | --- |
| ALLERGIC_RHINITIS | 25(3) |  | 23(2) | 12(4) |
| ASTHMA | 200(31) |  | 157(29) | 140(36) |
| CARD | 2(0) |  | 2(0) | 4(0) |
| DEPRESS | 2(0) |  | 1(0) | 0(0) |
| DYSLIPID | 166(0) |  | 120(0) | 91(0) |
| HEMORRHOIDS | 0 |  | 1(0) | 0 |
| HERNIA_ABDOMINOPELVIC | 2(1) |  | 2(1) | 1(1) |
| INSOMNIA | 0 |  | 0 | 0 |
| IRON_DEFICIENCY | 0 |  | 0 | 0 |
| IRRITABLE_BOWEL | 0 |  | 1(0) | 0 |
| MACDEGEN | 0 |  | 0 | 0 |
| OSTIOA | 1(0) |  | 2(0) | 2(1) |
| OSTIOP | 0 |  | 1(0) | 0 |
| PEPTIC_ULCERS | 1(0) |  | 1(0) | 0 |
| PSYCHIATRIC | 1 |  | 1 | 3 |
| PVD | 2(0) |  | 3(0) | 3(0) |
| STRESS | 1(0) |  | 1(0) | 1(0) |
| VARICOSE_VEINS | 40(2) |  | 22(2) | 20(2) |

#### LOAD results

After Bonferroni correction, TisCoMM-S^2^/S-MultiXcan/UTMOST identified 92/71/70 genome-wide significant genes, respectively, with 45 overlapping genes (17 of them are known LOAD GWAS genes). Here we define known LOAD GWAS gene as the ones reported in GWAS catalog. The qq-plots for associations in these three approaches are shown in Figure 5A. Among the 92 candidate target genes identified by TisCoMM-S^2^, 24 of them are previously known LOAD GWAS genes, which are annotated in the Manhattan plot in Figure 5A. These include genes on the chromosome (CHR) 2 (*BIN1*), CHR 6 (*CD2AP*), CHR 7 (*EPHA1*), CHR 8 (*CLU*), CHR 11 (*PICALM, CCDC89, MS4A2, MS4A6A*), CHR 16 (*IL34*), and CHR 19 (*STK11* and *APOE* region). Moreover, TisCoMM-S^2^ also identified 35 genes that were not significant in neither S-MultiXcan nor UTMOST, and four of them are known LOAD GWAS genes, including *IL34* (*p*-value =1 × 10^−6^), *PTK2b* (*p*-value =1.4 × 10^−9^), *EPHX* (*p*-value =4.7 × 10^−8^) and *STK11* (*p*-value = 7.2 × 10^−7^).

**Figure 5:**
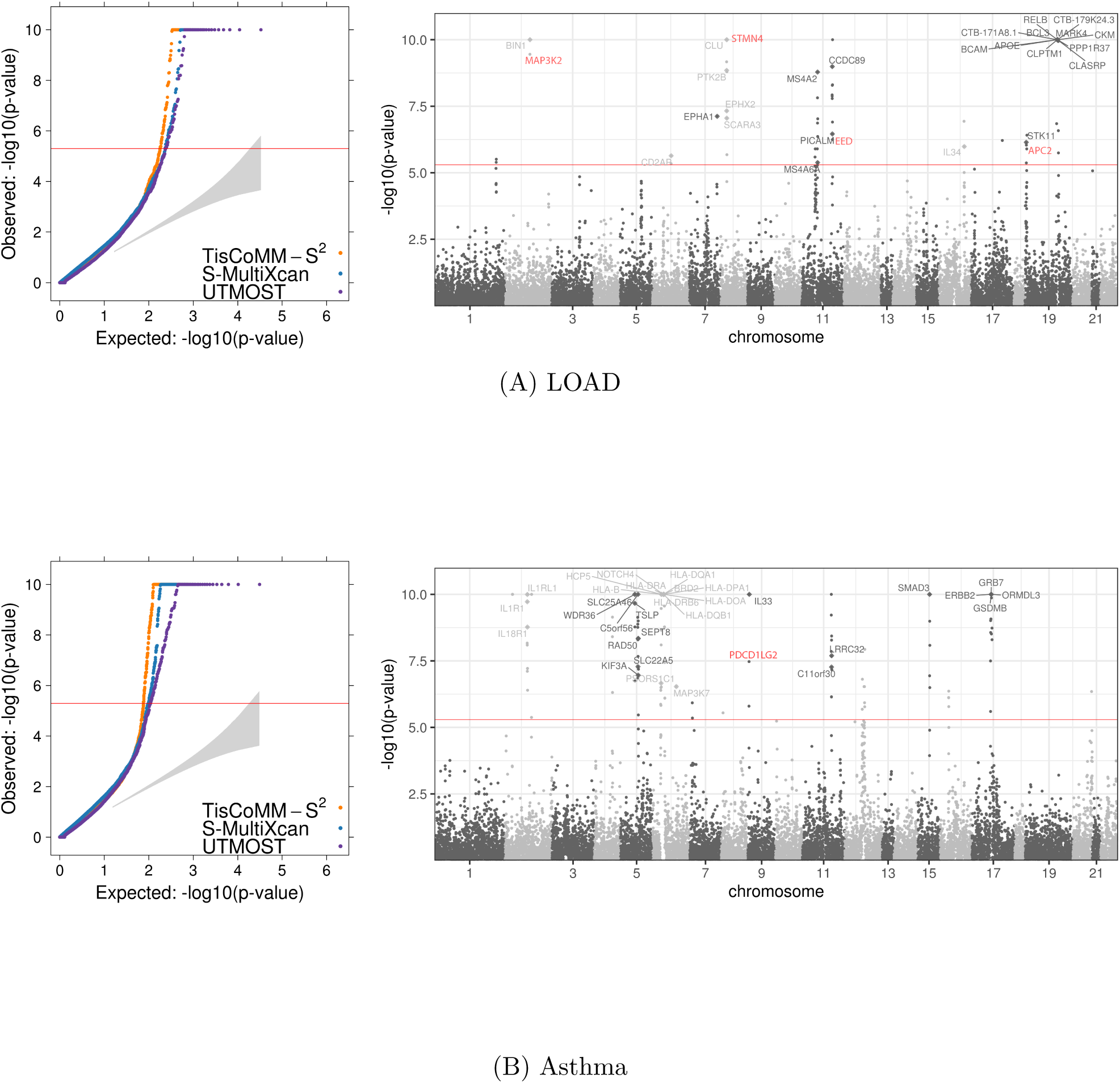
TisCoMM-S^2^ results for LOAD and asthma. The reference panel is European subsamples from 1000 Genome. In each row, the two panels show the qq-plot (left) and Manhatton plot (right).

Among all novel genes for LOAD identified by TisCoMM-S^2^, some of them were identified to be LOAD-related genes based on other computational models (e.g., *MAP3K2*) while some of them have not been directly linked to LOAD yet, but have been proven to be important regulators in different regions of the neuron system (e.g., *STMN4*, *EED* and *APC2*). *MAP3K2* is 200kb downstream of *B1N1*, a reported LOAD risk gene [31] that was also genome-wide significant in our joint test (*p*-values for both *B1N1* and *MAP3K2* < 10^−10^). *MAP3K2* belongs to the serine/threonine protein kinase family and has been previously identified as a member of the Alzheimer’s disease susceptibility network [32]. *STMN4* (*p*-value < 10^−10^) encodes the known protein that exhibits microtubule-destabilizing activity. The expression levels of this gene in mouse neurons have been shown to change significantly after different exposure of cortical nerve cells to the A*β* peptide [33]. The expression of *STMN4* in zebrafish has also been shown to have an important role in regulating neurogenesis in the neural keel stage [34]. *EED* (*p*-value =5.7 × 10^−7^) encodes a Polycomb protein, which plays a starring role as an important modulator of hippocampal development [35]. *APC2* (*p*-value = 1.3 × 10^−6^) is preferentially expressed in postmitotic neurons and involved in brain development through its regulation of neuronal migration and axon guidance [36]. We annotate these four genes in red in Figure 5A. Validation of these potential target genes requires further functional studies. The list of significant gene-trait associations of TisCoMM-S^2^, S-MultiXcan, and UTMOST can be found in Supplementary Table S2. To replicate our findings in another independent data set, we used the summary statistics from the GWAS by proxy (GWAX [37], the sample size is 114,564). Our replication rate was high (Supplementary Table S3), where 31 out of 92 genes were successfully replicated under the Bonferroni-corrected significance threshold and the numbers of replicated genes raised to 44 under a relaxed *p*-value cutoff of 0.05.

#### Asthma results

After Bonferroni correction, TisCoMM-S^2^/S-MultiXcan/UTMOST identified 200/157/140 genome-wide significant genes, respectively, with 98 overlapping genes in all three methods (and 21 of them are known asthma GWAS genes). The qq-plots for associations in these three approaches are shown in Figure 5B. Among all 200 candidate target genes identified by TisCoMM-S^2^, 31 of them are known asthma GWAS genes, which is annotated in the Manhattan plot in Figure 5B, including genes on CHR 2 (*IL1RL1/IL18R1*), CHR 5 (*TSLP/WDR36*, *RAD50*), CHR 6 (*HLA-DR/DQ* regions, *MAP3K7*), CHR 9 (*IL33*), CHR 11 (*C11orf30*, *LRRC32*), CHR 15 (*SMAD3*), and CHR 17 (genes from the 17q21 asthma locus). Also, TisCoMM-S^2^ identified 56 genes that were not significant in neither S-MultiXcan nor UTMOST, and two of them are known asthma GWAS genes, which are *PSORS1C1* (*p*-value =2.2 × 10^−7^), and *MAP3K7* (*p*-value =3 × 10^−7^).

Among all novel loci for asthma identified by TisCoMM-S^2^, *PDCD1LG2* was shown to have essential roles in modulating and polarizing T-cell functions in airway hyperreactivity [38]. Validating causal role of this gene in asthma requires further investigation. The list of significant gene-trait associations of TisCoMM-S^2^, S-MultiXcan, and UTMOST can be found in Supplementary Table S4. We annotate these two genes in red in Figure 5B.

To replicate our findings in another independent data set, we used the summary statistics from TAGC European-ancestry GWAS [39] (the sample size is 127,669). Our replication rate was high (Supplementary Table S5), where 179 out of 200 genes were successfully replicated under the Bonferroni-corrected significance threshold and the numbers of replicated genes raised to 189 under a relaxed *p*-value cutoff of 0.05.

### TisCoMM-S^2^ tissue-specific test infers gene effects in causal tissues

To demonstrate the utility of the TisCoMM-S^2^ tissue-specific test, we applied the tissue-specific test to all identified 92 candidate genes of LOAD and 200 candidate genes of asthma by using the TisCoMM-S^2^ joint test, and compared analysis results with those from CoMM [10, 11]. Table 3 shows the distributions of identified tissues with which candidate genes are associated in LOAD and asthma, respectively (see details in Supplementary Tables S6 and S7). Among all identified candidate genes respectively for both LOAD and asthma, 76.1% and 81.5% were significant in less than two tissues using TisCoMM-S^2^ while 70.7% and 60% were significant in all six tissues using CoMM-S^2^. The most plausible explanation is that compared to the multivariate perspective of our TisCoMM-S^2^ tissue-specific test, single-tissue approaches, e.g., CoMM-S^2^, tend to have larger tissue bias and more inflation in significant findings [9]. Suppose a gene is causal in tissue A but not in tissue B, and its expressions in tissues A and B are correlated. In a single-tissue test, the association can be spuriously significant for tissue B because of the similar gene expression pattern observed in both tissues. By performing a tissue-specific test for this gene in tissue B conditioned on tissue A, the significant spurious association will be largely excluded.

**Table 3:**
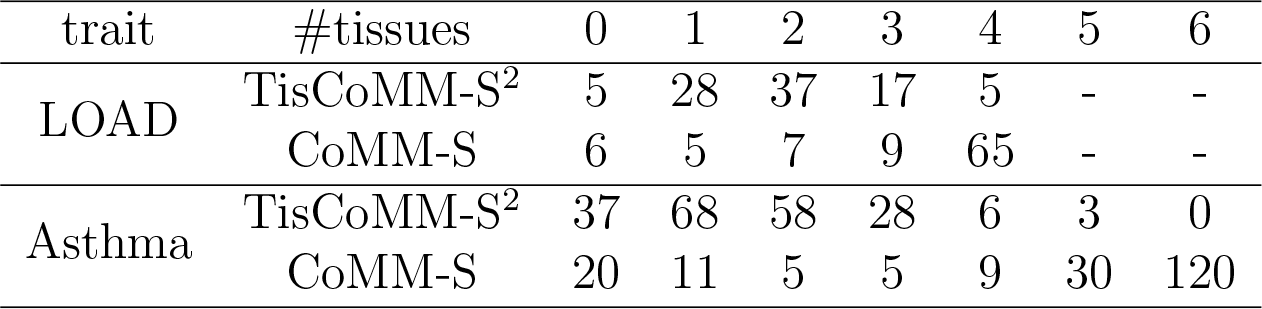
Distributions of tissues in which the candidate genes’ associations arise in LOAD and asthma.

To demonstrate the tissue-specific role of candidate genes inferred by TisCoM-S^2^ tissue-specific test for LOAD and asthma, respectively, we plot the volcano plots in Supplementary Figure S13, where the x-axis is the effect size showing in log scale, the y-axis is −*log*10 of the *p*-value from tissue-specific test, and the size of points reflect the cellular-heritability in each tissue. Known GWAS genes are also annotated. Next, we explored the tissue-specific effects of some well-replicated genes that are identified by the TisCoMM-S^2^ joint test for LOAD and asthma, respectively.

#### LOAD results

The well-replicated risk gene *APOE* [40] and its 50Kb downstream *CLPTM1* have been identified by the TisCoMM-S^2^ joint test. Moreover, the TisCoMM-S^2^ tissue-specific test identified *CLPTM1* to be significantly associated with LOAD in all four tissues (artery aorta, esophagus mucosa, nerve tibial, and skin sun-exposed lower leg with tissue-specific *p*-values < 4.9 × 10^−7^), but *APOE* to be only significantly associated with LOAD in artery aorta (tissue-specific *p*-value =8.3 × 10^−9^) and nerve tibial (tissue-specific *p*-value =1.2 × 10^−8^). On the other hand, CoMM-S^2^ significantly identified both *APOE* and *CLPTM1* in all four tissues (*p*-values ≤ 10^−10^) but failed to identify the difference of tissue-specific role for these two genes. We further investigate the molecular functions of LOAD associated genes in each tissue. In each of tested tissues in LOAD, there are about 40 tissue-specific genes. It is difficult to carry out a proper pathway analysis with such limited gene sets. So we classified the genes into seven functional groups based on which molecular functions they belong to. As shown in Figure 6A and 6B, majority (> 62%) of LOAD-associated genes belonged to binding and catalytic activity, and a small portion of significant LOAD genes were transcription factors suggesting that many regulation processes are going on at both protein and mRNA levels in different tissues.

**Figure 6:**
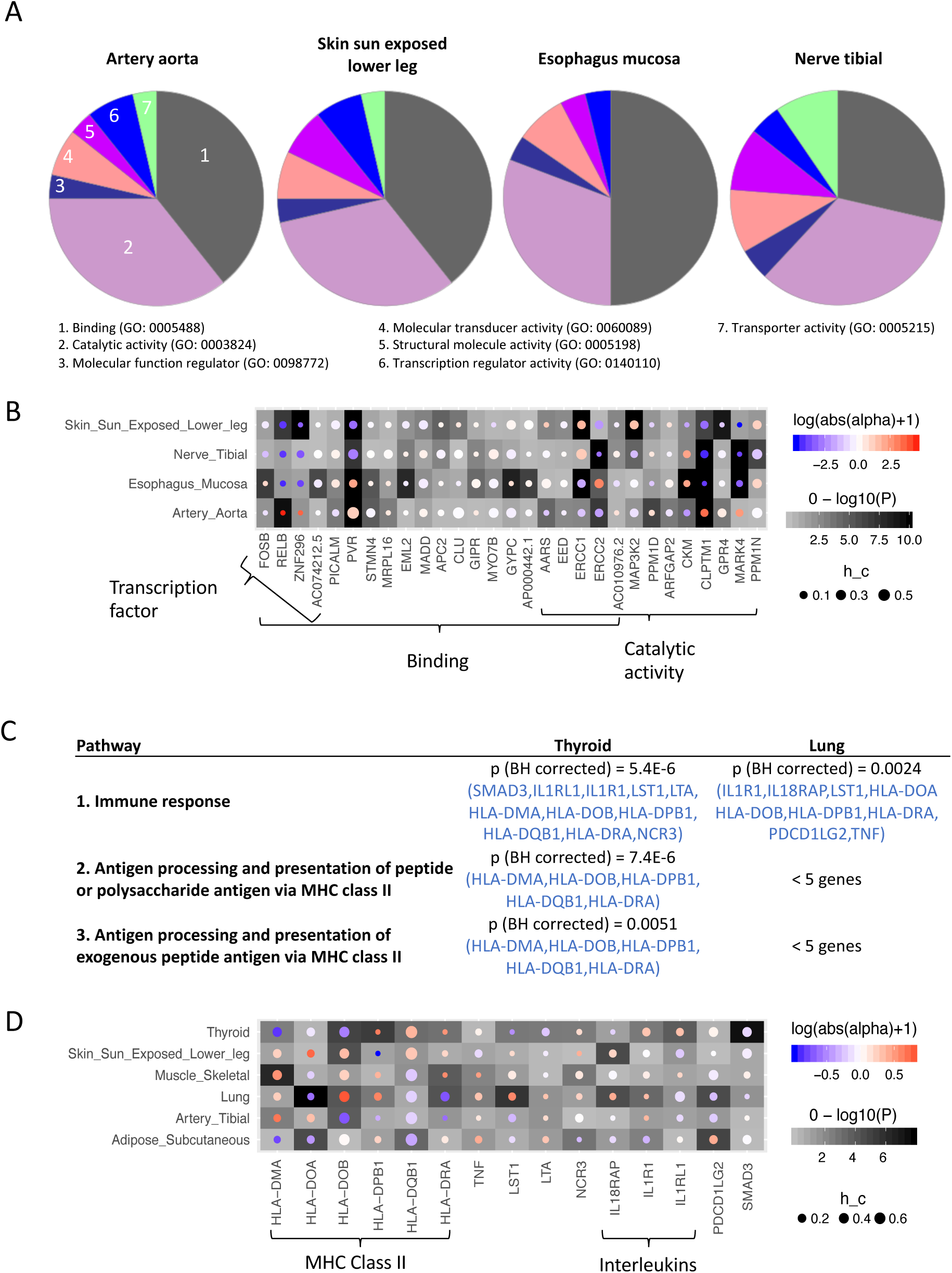
**A.** Each pie chart corresponding to a different tissue shows the percentage of LOAD-associated genes in each molecular function group (from gene ontology). **B.** The x-axis of the heatmap represents the union of LOAD-associated genes in 3 function groups (binding, catalytic activity, and transcription factor). The y-axis represents different tissue types. In each cell, the background color (shades of gray) indicates the significance level, the circle size indicates the heritability, and the color inside each circle indicates the effect size. **C.** Pathway analysis of asthma-associated genes in thyroid and lung. Pathway analysis was done using a web-based software DAVID, testing the enrichments of asthma-associated genes in biological processes (from gene ontology). Significant pathways were selected if gene count ≥ 5 and Benjamini-Hochberg (BH) corrected p-value ≤ 0.05. The asthma-associated genes are highlighted in blue. **D.** The x-axis of the heatmap represents the asthma-associated genes in the immune response pathway. And all the other settings are the same as the one used in part B.

According to our tissue selection strategy, above tissue-specific test for LOAD was conducted on four non-brain tissues (enriched tissues). To further investigate the gene expression changes in the well-studied disease tissues, three more brain regions (hippocampus, frontal cortex, and cerebellar hemisphere) were selected for another tissue-specific analysis for LOAD. Because it is known that hippocampus is one of the first brain regions to be affected by Alzheimer’s disease and related to the memory lost [41], markers such as A*β* in frontal cortex can be used to predict future Alzheimer’s disease [42], and cerebellum is affected in the final stage of the disease and related to cognitive decline [43]. The joint test conducted on brain regions revealed 105 LOAD associated genes, of which 73 were identified in the enriched tissues (Figure S14A), and the other 32 genes were uniquely identified in brain regions (Figure S14B). The most significant gene uniquely identified in brain regions is *KLC3* according to the joint test (*p*-value < 10^−10^), which is within 50kb downstream of *APOE*. Moreover, it is significantly associated with LOAD in hippocampus region only, but not the other two brain regions according to the tissue-specific test (Figure S14B). Thus, we propose *KLC3* as one of the potential novel targets for LOAD in hippocampus.

#### Asthma results

We take identified genes ORMDL3 and GSDMB in the 17q21 asthma locus as an example, because these two genes have been mentioned as asthma susceptibility locus by many studies, a comprehensive review was written by Stein et al. [44]. The original finding of ORMDL3 was observed in one GWAS study, and have been further validated in a mouse model [45]. The TisCoMM-S^2^ tissue-specific test identified both *ORMDL3* and *GSDMB* to be significantly associated with asthma only in lung tissue (see the volcano plot in Supplementary Figure S14B, tissue-specific *p*-values for these two genes are 1.7 × 10^−3^ and 7.1 × 10^−7^, respectively). However, CoMM-S^2^ identified both *ORMDL3* and *GSDMB* in all six tissues (*p*-values ≤ 10^−10^) but failed to identify the relevant tissues with which these two genes are causally related to asthma. We further conducted pathway analysis using DAVID [46] on six sets of asthma-associated genes in all six tissues (thyroid, lung, artery tibial, muscle skeletal, adipose subcutaneous, and skin sun-exposed lower leg), respectively. As listed in Figure 6B, all three significant pathways in thyroid tissue belonged to the immune system, and the only significant pathway in lung tissue was immune response. However, no significant pathways were detected in the other four tissues. Among asthma-associated genes in immune response (first row in Figure 6C and 6D), the majority of them were shared between thyroid and lung, and located in the MHC region on CHR 6 including several *HLA* genes and *LST1*. Our pathway analysis suggests that nearly the same set of immune genes in thyroid and lung are responsible for asthma development.

## Discussion

Despite the substantial successes of TWAS and its variants, the existing multi-tissue methods have several limitations, e.g., incapability to identify the tissue-specific effect of a gene, ignorance of imputation uncertainty, and failure to efficiently use tissue-shared patterns in eQTLs. To overcome these limitations and provide additional perspectives over tissue-specific roles of identified genes, we have proposed a powerful multi-tissue TWAS model, together with a computationally efficient inference method and software implementation in TisCoMM. Specifically, we have developed a joint test for prioritizing gene-trait associations and a tissue-specific test for identifying the tissue-specific role of candidate genes. Conditioned on the inclusion of trait-relevant tissues, the tissue-specific test in TisCoMM can mostly remove the spurious associations in a single-tissue test due to high correlations among gene expression across tissues. We have also developed a summary-statistic-based model, TisCoMM-S^2^, extending the applicability of TisCoMM to publicly available GWAS summary data. Using both simulations and real data, we examined the relationship between TisCoMM and TisCoMM-S^2^. Our results, as shown in Figure 2, show that the test statistics from TisCoMM and TisCoMM-S^2^ are highly correlated (*R*^2^ > 0.95). We further analyzed summary-level GWAS data from 33 traits with replication data for Alzheimer’s disease and asthma. Overall, the findings from TisCoMM-S^2^ are around 30% more than those from S-MultiXcan or UTMOST while qq-plots from these studies show that there are no apparent inflations. To replicate our findings for Alzheimer’s disease and asthma, we applied TisCoMM-S^2^ to independent data sets for each disease. Results show that replication rates for Alzheimer’s disease and asthma are high.

We further inferred the tissue-specific effects of identified genes using the TisCoMM-S^2^ tissue-specific test. By classifying these genes into seven functional groups, we observed that majority (62%) of LOAD-associated genes were related to binding and catalytic activity while a small portion was from transcription factors suggesting active regulation processes at both protein and mRNA level in different tissues. We also observed about 40 LOAD-associated genes in each non-brain tissues. The significance of these genes could be due to the exclusion of LOAD-relevant tissues, e.g., brain tissues. To fill this gap, we further conducted one more analysis on three brain regions, and identified 32 brain specific genes. For asthma, genes *ORMDL3* and *GSDMB* were identified to be significantly associated with asthma only in lung tissue using TisCoMM-S^2^ tissue-specific test. However, single-tissue analysis (CoMM-S^2^) identified both genes significant in all six tested tissues. Further pathway analysis shows that all three significant pathways for thyroid tissue belong to the immune system and the only significant pathway for lung tissue was immune response. The majority of shared genes between thyroid and lung tissues are located in the MHC region on CHR 6, including several *HLA* genes and *LST1*. The proteins encoded by *HLA* genes are known as antigens. In combination with antigen-presenting cells (e.g., macrophages and dendritic cells), they play an essential role in the activation of immune cells as well as airway inflammation in response to asthma-related allergens [47, 48]. Based on our tissue-specific test, *TNF* that is a well-studied asthma gene [49, 50] was explicitly identified to be associated with asthma in lung tissue. The positive correlation between *TNF* expression and asthma in lung confirmed our previous understanding of *TNF* activation in asthma, promoting airway inflammation and airway hyperresponsiveness. On the other hand, *LTA* was specifically regulated in thyroid tissue. It is a cytokine produced by lymphocytes, and also known as a regulator of lipid metabolism [51]. Another immune gene regulated individually in thyroid tissue is *NCR3*, which mediates the crosstalk between natural killer cells and dendritic cells [52]. However, it remains unclear how the alteration of *LTA* and *NCR3* in thyroid could lead to asthma development.

Despite the utility of TisCoMM to perform gene-trait association analysis in a tissue-specific manner, it is primarily designed to test genes with direct effects from cis-eQTL. Recently, an omnigenic model was proposed to better understand the underlying mechanism of so-called polygenicity in complex traits [53]. Liu et al. [54] further provided a theoretical model to understand complex trait architecture by partitioning genetic contributions into direct effects from core genes and indirect effects from peripheral genes acting in trans. Most works from TWAS identify core genes with direct effects. How to effectively interrogate peripheral genes with indirect effects essentially remains an open question. As high-throughput data are continuously generating for a much larger sample size with more precision, TisCoMM sheds light on how to integrate useful data for the desired analysis effectively.

## Methods

### Model settings

Conventionally, both single-tissue and multi-tissue TWAS methods proceed by conducting a prediction model in Equation (1) followed by a subsequent association analysis in Equation (2), where a steady-state gene expression is imputed from 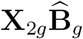 and 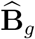 is estimated in the first prediction model, e.g., PrediXcan, MultiXcan, S-MulitXcan, and UTMOST. However, this imputation strategy ignores the uncertainty in the process of expression imputation. Here, we describe the individual-level data version of TisCoMM by jointly analyzing models (1) and (2), and extensions to summary statistics will be discussed in the Supplementary Text. Assume 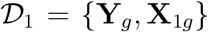 denote the reference transcriptome data set of gene *g* for *n*_1_ samples over *T* tissues, where **Y**_*g*_ is the *n*_1_ × *T* expression matrix for this gene over *T* tissues, and **X**_1*g*_ is the corresponding *n*_1_ × *M*_*g*_ standardized genotype matrix for *M*_*g*_ cis-SNPs within this gene. Denote the GWAS data 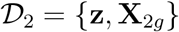, where **z** is an *n*_2_ × 1 vector of phenotypic values, **X**_2*g*_ is the corresponding *n*_2_ × *M*_*g*_ standardized genotype matrix for *M*_*g*_ cis-SNPs. Since we conduct hypothesis testing sequentially or parralelly for each gene, we will omit the subscript *g* in all the expression that has dependence on gene *g* to simplify notations. Our model becomes

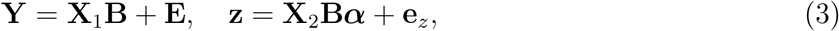

where ***α*** ∈ ℝ^*T*^, 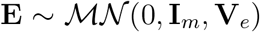, and 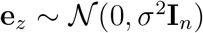. Note that we assume 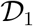 and 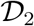 are centered and thus intercepts can be omitted.

To estimate the tissue-specific eQTL effects, we need to first estimate an *M* × *T* coefficient matrix **B**. To reduce the number of parameters, we follow an adaptive weighting scheme [22, 23, 24]: we regress the gene expression in tissue type *t* on the *j*th eQTL and let the marginal eQTL effect be the adaptive weight, *w*_*jt*_. Specifically, we assume the joint eQTL effect size *β*_*jt*_ can be decomposed into variant-dependent components *b*_*j*_ and tissue-specific components *w*_*jt*_: *β*_*jt*_ = *b_j_w*_*jt*_. That is, **B** = diag{**b**}**W**. Similar strategies have been applied to model tissue-shared patterns [24, 21]. Let **y**_*i*_, **x**_1*i*_ and **w**_*j*_ denote the *i*th row of **Y**, **X**_1_ and **W**, respectively. Our model can be written as

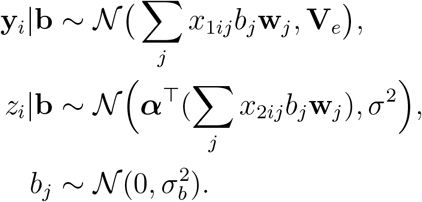

Denote ***θ*** = (***α***, 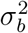, *σ*^2^, **V**_*e*_)^*T*^ the vector for all model parameters. We need to estimate parameters and maker inference for ***α***. Both the TisCoMM joint test and tissue-specific test are based on likelihood ratio tests. The joint test for gene-trait associations can be formally set up as *H*_0_ : ***α*** = 0 verses *H*_1_ : ***α*** ≠ 0. The corresponding likelihood ratio test statistic is given by

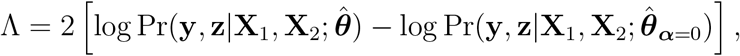

where 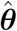 is the vector of parameter estimates under the full model, and 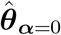 is the vector of estimates under the constrain ***α*** = 0. Similarly, the tissue-specific test for the tissue-specific effect can be formally set up as *H*_0_ : *α*_*t*_ = 0 verses *H*_1_ : *α*_*t*_ ≠ 0. The corresponding likelihood ratio test statistic is given by

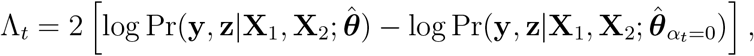

where 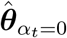 is the vector of parameter estimates under *α*_*t*_ = 0.

For statistical inference, we developed an expectation-maximization (EM) algorithm accelerated by expanding parameters [55]. Details of updating equations for each parameter and the corresponding algorithm can be found in Supplementary Text.

### GWAS data

#### The NFBC1966 data set

The NFBC1966 data set consists of ten traits and 364,590 SNPs from 5402 individuals [26], including total cholesterol (TC), high-density lipoprotein cholesterol (HDL-C), low-density lipoprotein cholesterol (LDL-C) and triglycerides (TG), inflammatory marker C-reactive protein, markers of glucose homeostasis (glucose and insulin), body mass index (BMI) and blood pressure (BP) measurements (systolic and diastolic BP). Quality control procedures are conducted following similar steps to Shi et al. [56]. Specifically, individuals with missing-ness in any of the traits and with genotype missing call-rates > 5% were excluded. We excluded SNPs with minor allele frequency (MAF) < 1%, missing call-rates > 1%, or failed Hardy-Weinberg equilibrium. After quality control filtering, 172,412 SNPs from 5123 individuals were available for downstream analysis.

The tissues used in TisCoMM and TisCoMM-S^2^ were the same, and the six tissues with the largest number of overlapped individuals were used. The summary statistics for TisCoMM-S^2^ were calculated using PLINK [57].

#### Summary-level GWAS data

We obtained summary statistics from GWASs for 33 traits, including 15 traits from [19] and 18 traits from the UK Biobank. Details of these traits can be found in Supplementary Table S1. In the main text, we discussed LOAD and asthma. Analyses results for other traits can be found in Supplementary Text.

### GTEx eQTL Data

Th GTEx data including genotype and RNA-seq data are obtained from dbGaP with accession number phs000424.v7.p2. Processed gene-expression data are available on the GTEx portal (https://gtexportal.org/home/). In the eQTL data, we removed SNPs with ambiguous alleles or MAF less 0.01.

We used two different strategies to select tissues used in our real data analysis. For the 15 NG traits, we obtained the top enriched tissues for each trait according to Supplementary Table 2 in [19], and a subset of tissues with sample sizes larger than 100 was kept. For the UKB traits, we used the six tissues with the largest number of overlapped individuals.

### Reference panel

Due to the absence of genotype data using summary statistics, we use reference samples to estimate the LD structures *R* among SNPs in the study samples. Since diseases and traits considered in our real data application are for European population cohorts, we choose to use European subsamples from the 1000 Genome Project as a reference panel.

Let **X**_*r*_ denote the genotype matrix for cis-SNPs in the reference panel. To estimate the LD matrix *R*, we adopt a simple shrinkage method as follows. We first calculate the empirical correlation matrix 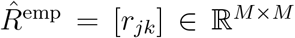 with 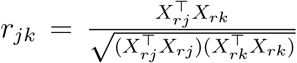 where *X*_*rj*_ the *j*th column of **X**_*r*_. To make the estimated correlation matrix positive definite, we apply a simple shrinkage estimator [58]: 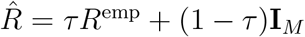, where *τ* ∈ [0, 1] is the shrinkage intensity. In real data application, we fixed the shrinkage intensity at 0.95 both for simplicity and computational stability.

### Web Resources

*TisCoMM* is available at https://github.com/XingjieShi/TisCoMM/.

*PrediXcan, MultiXcan* and *S-MultiXcan* are available at http://gene2pheno.org/.

*UTMOST* is available at https://github.com/Joker-Jerome/UTMOST/.

*CoMM* is available at https://github.com/gordonliu810822/CoMM.

Known trait-associated genes are available at the NHGRI-EBI GWAS Catalog https://www.ebi.ac.uk/gwas/.

Summary statistics from UK Biobank is available at http://geneatlas.roslin.ed.ac.uk/.

URLs for summary statistics from Gamazon et al. [19] are summarized in Supplementary Table S1.

## Acknowledgements

This work was supported by grant R-913-200-098-263 from the Duke-NUS Medical School, AcRF Tier 2 (MOE2016-T2-2-029, MOE2018-T2-1-046 and MOE2018-T2-2-006) from the Ministry of Education, Singapore, grant No. 71501089, No. 11501579 and No. 71472023 from the National Natural Science Foundation of China; and grant Nos. 22302815, No. 12316116 and No. 12301417 from the Hong Kong Research Grant Council. The computational work for this article was partially performed using resources from the National Supercomputing Centre, Singapore (https://www.nscc.sg).

