## Supplementary Text for "A tissue-specific collaborative mixed model for jointly analyzing multiple tissues in transcriptome-wide association studies"

**3 Additional results for NFBC1966 study** **10**

**4 Additional results for simulations** **12**

**5 Additional results for NG and UKB traits** **18**

### 1 Statistical Model for TisCoMM

#### 1.1 Model settings

We describe the individual data version of TisCoMM in this section, and extensions to summary statistics will discussed in next section. Assume $\mathcal{D}_1 = \{\mathbf{Y}_g, \mathbf{X}_{1g}\}$ denote the reference transcriptome data set of gene $g$ for $n_1$ samples over $T$ tissues, e.g. $\mathbf{Y}_g \in \mathbb{R}^{n_1 \times T}$ is the expression matrix for this gene over $T$ tissues, $\mathbf{X}_{1g} \in \mathbb{R}^{n_1 \times M_g}$ is the genotype matrix for cis-SNPs within this gene. Denote the GWAS data $\mathcal{D}_2 = \{\mathbf{z}, \mathbf{X}_{2g}\}$, where $\mathbf{z}$ is an $n_2 \times 1$ vector of phenotypic values, $\mathbf{X}_{2g}$ is the genotype matrix for $M_g$ cis-SNPs.

To simplify notation we will omit the subscript $g$ in all the expression that has dependence on gene $g$. Our model is

$$\mathbf{Y} = \mathbf{X}_1\mathbf{B} + \mathbf{E}, \tag{S1}$$

$$\mathbf{z} = \mathbf{X}_2\mathbf{B}\boldsymbol{\alpha} + \mathbf{e}_z, \tag{S2}$$

where $\boldsymbol{\alpha} \in \mathbb{R}^T$, $\mathbf{E} \sim \mathcal{MN}(0, \mathbf{I}_m, \mathbf{V}_e)$, and $\mathbf{e}_z \sim \mathcal{N}(0, \sigma^2\mathbf{I}_n)$. Note that we assume $\mathcal{D}_1$ and $\mathcal{D}_2$ are centered and thus intercepts can be omitted.

To estimate the tissue-specific eQTL effects, we need to estimate a $M \times T$ coefficient matrix $\mathbf{B}$. To reduce the number of parameters, we follow an adaptive weighting scheme [1, 2, 3]: we regress the gene expression in tissue type $t$ on the $j$th eQTL and let the marginal eQTL effect be the adaptive weight, $w_{jt}$. Specifically, we assume the joint eQTL effect size $\beta_{jt}$ can be decomposed into SNP-dependent components $b_j$ and marginal tissue-specific effect size $w_{jt}$: $\beta_{jt} = b_j w_{jt}$. That is, $\mathbf{B} = \text{diag}\{\mathbf{b}\}\mathbf{W}$. In the GTEx data, empirical evidence is observed supporting the validity of this assumption [3].

Let $\mathbf{y}_i$, $\mathbf{x}_{1i}$ and $\mathbf{w}_j$ denote the $i$th row of $\mathbf{Y}$, $\mathbf{X}_1$ and $\mathbf{W}$, respectively. Our model is

$$\begin{aligned} \mathbf{y}_i|\mathbf{b} &\sim \mathcal{N}\big(\sum_j x_{1ij} b_j \mathbf{w}_j, \mathbf{V}_e\big), \\ z_i|\mathbf{b} &\sim \mathcal{N}\Big(\boldsymbol{\alpha}^\top(\sum_j x_{2ij} b_j \mathbf{w}_j), \sigma^2\Big), \\ b_j &\sim \mathcal{N}(0, \sigma_b^2). \end{aligned} \tag{S3}$$

Let $\boldsymbol{\theta} = (\boldsymbol{\alpha}, \sigma_b^2, \sigma^2, \mathbf{V}_e)$ denote all model parameters. We need to estimate parameters and do inference for $\boldsymbol{\alpha}$. Both TisCoMM joint test and tissue-specific test are conducted by likelihood ratio test. The joint test for gene-trait associations is $H_0 : \boldsymbol{\alpha} = 0$ verses $H_1 : \boldsymbol{\alpha} \neq 0$. The likelihood ratio test statistic is given by

$$\Lambda = 2\left[\log \Pr(\mathbf{y}, \mathbf{z}|\mathbf{X}_1, \mathbf{X}_2; \hat{\boldsymbol{\theta}}) - \log \Pr(\mathbf{y}, \mathbf{z}|\mathbf{X}_1, \mathbf{X}_2; \hat{\boldsymbol{\theta}}_{\boldsymbol{\alpha}=0})\right],$$

where $\hat{\boldsymbol{\theta}}$ is the parameter estimator, and $\hat{\boldsymbol{\theta}}_{\boldsymbol{\alpha}=0}$ is the estimator under constrain $\boldsymbol{\alpha} = 0$. Similarly, the tissue-specific test for tissue specific effects is $H_0 : \alpha_t = 0$ verses $H_1 : \alpha_t \neq 0$. The likelihood ratio test statistic is given by

$$\Lambda_t = 2\left[\log \Pr(\mathbf{y}, \mathbf{z}|\mathbf{X}_1, \mathbf{X}_2; \hat{\boldsymbol{\theta}}) - \log \Pr(\mathbf{y}, \mathbf{z}|\mathbf{X}_1, \mathbf{X}_2; \hat{\boldsymbol{\theta}}_{\alpha_t=0})\right],$$

where $\hat{\boldsymbol{\theta}}_{\alpha_t=0}$ is the parameter estimator under $\alpha_t = 0$.

#### 1.2 Estimation procedure

For statistical inference, we developed an expectation-maximization (EM) algorithm, and adopted parameter expansion technique [4] to accelerate the convergence rate of standard EM algorithm. Specifically, we expanded our model as follows:

$$\mathbf{Y} = \lambda \mathbf{X}_1 \mathbf{B} + \mathbf{E}, \tag{S4}$$

$$\mathbf{z} = \mathbf{X}_2 \mathbf{B} \boldsymbol{\alpha} + \mathbf{e}_z, \tag{S5}$$

where $\lambda \in \mathbb{R}$ is the expanded parameter and our expanded model parameter become $\boldsymbol{\theta} = (\boldsymbol{\alpha}, \sigma_b^2, \sigma^2, \mathbf{V}_e, \lambda)$.

Let $\mathbf{D}_{1i} = \mathbf{W}^\top \text{diag}\{\mathbf{x}_{1i}\}$ and $\mathbf{D}_{2i} = \mathbf{W}^\top \text{diag}\{\mathbf{x}_{2i}\}$. The complete data log-likelihood can be written as

$$\begin{aligned}
&\log \Pr(\mathbf{y}, \mathbf{z}, \mathbf{b} | \mathbf{X}_1, \mathbf{X}_2; \boldsymbol{\theta}) \\
=& -\frac{n_1 T}{2} \log 2\pi - \frac{n_1}{2} \log |\mathbf{V}_e| - \frac{1}{2} \sum_{i=1}^{n_1} (\mathbf{y}_i - \lambda \mathbf{D}_{1i} \mathbf{b})^\top \mathbf{V}_e^{-1} (\mathbf{y}_i - \lambda \mathbf{D}_{1i} \mathbf{b}) \\
& - \frac{n_2}{2} \log 2\pi - \frac{n_2}{2} \log \sigma^2 - \frac{1}{2\sigma^2} \sum_{i=1}^{n_2} (z_i - \boldsymbol{\alpha}^\top \mathbf{D}_{2i} \mathbf{b})^2 \\
& - \frac{M}{2} \log 2\pi - \frac{M}{2} \log \sigma_b^2 - \frac{1}{2\sigma_b^2} \mathbf{b}^\top \mathbf{b}.
\end{aligned} \tag{S6}$$

By checking the term regarding $\mathbf{b}$ in the complete data log-likelihood, we would know the posterior of $\mathbf{b}$ should be of Gaussian distribution $\mathcal{N}(\mathbf{b}|\boldsymbol{\mu}, \boldsymbol{\Sigma})$ with

$$\begin{aligned}
\boldsymbol{\Sigma}^{-1} =& \lambda^2 \sum_{i=1}^{n_1} \mathbf{D}_{1i}^\top \mathbf{V}_e^{-1} \mathbf{D}_{1i} + \sum_{i=1}^{n_2} \mathbf{D}_{2i}^\top \boldsymbol{\alpha} \boldsymbol{\alpha}^\top \mathbf{D}_{2i} / \sigma^2 + \frac{1}{\sigma_b^2} \mathbf{I}_M, \\
\boldsymbol{\mu} =& \boldsymbol{\Sigma} \left[ \lambda \sum_{i=1}^{n_1} \mathbf{D}_{1i}^\top \mathbf{V}_e^{-1} \mathbf{y}_i + \sum_{i=1}^{n_2} \mathbf{D}_{2i}^\top z_i \boldsymbol{\alpha} / \sigma^2 \right].
\end{aligned} \tag{S7}$$

#### E-step

Let $\boldsymbol{\theta}^{old}$ be the current estimate of all parameters. To simplify notation, let

$$\begin{aligned}
\hat{\mathbf{B}} &= \text{diag}\{\boldsymbol{\mu}\} \mathbf{W} \\
\boldsymbol{\Sigma}_i &= \mathbf{D}_{1i} \boldsymbol{\Sigma} \mathbf{D}_{1i}^\top, \\
\mathbf{r}_i &= \mathbf{y}_i - \lambda \mathbf{D}_{1i} \boldsymbol{\mu}.
\end{aligned} \tag{S8}$$

Taking expectation of complete data log-likelihood with respect to the current conditional distribution of $\mathbf{b}$ given the current estimates of the parameters $\boldsymbol{\theta}^{old}$, we have

$$
\begin{aligned}
& Q(\boldsymbol{\theta}|\boldsymbol{\theta}^{old}) \\
=& \mathbb{E}\left[\log \Pr(\mathbf{y}, \mathbf{z}, \mathbf{b}|\mathbf{X}_1, \mathbf{X}_2; \boldsymbol{\theta})|\boldsymbol{\theta}^{old}\right] \\
=& -\frac{n_1 T}{2}\log 2\pi - \frac{n_1}{2}\log|\mathbf{V}_e| - \frac{1}{2}\sum_{i=1}^{n_1}\left[\mathbf{r}_i^\top \mathbf{V}_e^{-1}\mathbf{r}_i + \lambda^2 \mathrm{Tr}(\mathbf{V}_e^{-1}\boldsymbol{\Sigma}_i)\right] \\
& - \frac{n_2}{2}\log 2\pi - \frac{n_2}{2}\log \sigma^2 - \frac{1}{2\sigma^2}\sum_{i=1}^{n_2}\left[(z_i - \boldsymbol{\alpha}^\top \mathbf{D}_{2i}\boldsymbol{\mu})^2 + \mathrm{Tr}(\boldsymbol{\alpha}^T \mathbf{D}_{2i}\boldsymbol{\Sigma}\mathbf{D}_{2i}^\top \boldsymbol{\alpha})\right] \\
& - \frac{M}{2}\log 2\pi - \frac{M}{2}\log \sigma_b^2 - \frac{1}{2\sigma_b^2}\left[\boldsymbol{\mu}^\top \boldsymbol{\mu} + \mathrm{Tr}(\boldsymbol{\Sigma})\right].
\end{aligned} \tag{S9}
$$

#### M-step

By setting the first partial derivative of $Q$ to zero, we obtained the updates for all parameters as follows:

$$
\begin{aligned}
\mathbf{V}_e &= \frac{1}{n_1}\sum_{i=1}^{n_1}(\mathbf{r}_i\mathbf{r}_i^\top + \lambda^2 \boldsymbol{\Sigma}_i), \\
\sigma_b^2 &= \frac{1}{M}\left[\boldsymbol{\mu}^\top \boldsymbol{\mu} + \mathrm{Tr}(\boldsymbol{\Sigma})\right], \\
\sigma^2 &= \frac{1}{n_2}\sum_{i=1}^{n_2}\left[(z_i - \boldsymbol{\alpha}^\top \mathbf{D}_{2i}\boldsymbol{\mu})^2 + \mathrm{Tr}(\boldsymbol{\alpha}^\top \mathbf{D}_{2i}\boldsymbol{\Sigma}\mathbf{D}_{2i}^\top \boldsymbol{\alpha})\right], \\
\boldsymbol{\alpha} &= \left[\sum_{i=1}^{n_2}\left(\mathbf{D}_{2i}\boldsymbol{\mu}\boldsymbol{\mu}^\top \mathbf{D}_{2i}^\top + \mathbf{D}_{2i}\boldsymbol{\Sigma}\mathbf{D}_{2i}^\top\right)\right]^{-1}\sum_{i=1}^{n_2} z_i \mathbf{D}_{2i}\boldsymbol{\mu}, \\
\lambda &= \sum_{i=1}^{n_1}\boldsymbol{\mu}^T \mathbf{D}_{1i}^\top \mathbf{V}_e^{-1}(\mathbf{y}_i - \boldsymbol{\eta}^\top \mathbf{c}_{1i})\left[\sum_{i=1}^{n_1}\boldsymbol{\mu}^\top \mathbf{D}_{1i}^\top \mathbf{V}_e^{-1}\mathbf{D}_{1i}\boldsymbol{\mu} + \sum_{i=1}^{n_1}\mathrm{Tr}(\mathbf{V}_e^{-1}\Sigma_i)\right]^{-1}.
\end{aligned} \tag{S10}
$$

#### Reduction-step

We reset the estimation of parameters using the reduction function:

$$
R(\boldsymbol{\alpha}, \sigma_b^2, \sigma^2, \mathbf{V}_e, \lambda) = (\boldsymbol{\alpha}/\lambda, \lambda^2\sigma_b^2, \sigma^2, \mathbf{V}_e). \tag{S11}
$$

#### 1.3 Likelihood Computation

To perform likelihood ratio test, we need to compute the observed likelihood function $\log \Pr(\mathbf{y}, \mathbf{z}|\mathbf{X}_1, \mathbf{X}_2; \hat{\boldsymbol{\theta}})$ efficiently. We note that

$$\begin{aligned}\log \Pr(\mathbf{y}, \mathbf{z}|\mathbf{X}_1, \mathbf{X}_2; \hat{\boldsymbol{\theta}}) &= \mathbb{E} \log \Pr(\mathbf{y}, \mathbf{z}|\mathbf{X}_1, \mathbf{X}_2; \hat{\boldsymbol{\theta}}) \\ &= \mathbb{E} \left[\log \Pr(\mathbf{y}, \mathbf{z}, \mathbf{b}|\mathbf{X}_1, \mathbf{X}_2; \boldsymbol{\theta})\right] - \mathbb{E} \log \Pr(\mathbf{b}|\mathbf{y}, \mathbf{z}, \mathbf{X}_1, \mathbf{X}_2; \boldsymbol{\theta}),\end{aligned} \tag{S12}$$

where the first and second terms in the last equation are $Q$ function (S9) and entropy of a multivariate normal distribution respectively. Both of them can be easily calculated.

#### 1.4 EM Algorithm

Implementation details for our EM algorithm with parameter expansion (PXEM) are summarized in Algorithm 1 for clarity.

**Algorithm 1:** PXEM

Initialize $\boldsymbol{\theta} = (\boldsymbol{\alpha}, \sigma_b^2, \sigma^2, \mathbf{V}_e, \lambda)$;

**repeat**

**E-step:**

$$\boldsymbol{\Sigma}^{-1} = \lambda^2 \sum_{i=1}^{n_1} \mathbf{D}_{1i}^\top \mathbf{V}_e^{-1} \mathbf{D}_{1i} + \sum_{i=1}^{n_2} \mathbf{D}_{2i}^\top \boldsymbol{\alpha}\boldsymbol{\alpha}^\top \mathbf{D}_{2i}/\sigma^2 + \frac{1}{\sigma_b^2}\mathbf{I}_M,$$

$$\boldsymbol{\mu} = \boldsymbol{\Sigma}\left[\lambda \sum_{i=1}^{n_1} \mathbf{D}_{1i}^\top \mathbf{V}_e^{-1}\mathbf{y}_i + \sum_{i=1}^{n_2} \mathbf{D}_{2i}^\top z_i \boldsymbol{\alpha}/\sigma^2\right];$$

**M-step:**

$$\mathbf{V}_e = \frac{1}{n_1}\sum_{i=1}^{n_1}(\mathbf{r}_i\mathbf{r}_i^\top + \lambda^2\boldsymbol{\Sigma}_i),$$

$$\sigma_b^2 = \frac{1}{M}\left[\boldsymbol{\mu}^\top\boldsymbol{\mu} + \mathrm{Tr}(\boldsymbol{\Sigma})\right],$$

$$\sigma^2 = \frac{1}{n_2}\sum_{i=1}^{n_2}\left[(z_i - \boldsymbol{\alpha}^\top \mathbf{D}_{2i}\boldsymbol{\mu})^2 + \mathrm{Tr}(\boldsymbol{\alpha}^\top \mathbf{D}_{2i}\boldsymbol{\Sigma}\mathbf{D}_{2i}^\top\boldsymbol{\alpha})\right],$$

$$\boldsymbol{\alpha} = \left[\sum_{i=1}^{n_2}\left(\mathbf{D}_{2i}\boldsymbol{\mu}\boldsymbol{\mu}^\top\mathbf{D}_{2i}^\top + \mathbf{D}_{2i}\boldsymbol{\Sigma}\mathbf{D}_{2i}^\top\right)\right]^{-1}\sum_{i=1}^{n_2} z_i\mathbf{D}_{2i}\boldsymbol{\mu},$$

$$\lambda = \sum_{i=1}^{n_1}\boldsymbol{\mu}^T\mathbf{D}_{1i}^\top\mathbf{V}_e^{-1}(\mathbf{y}_i - \boldsymbol{\eta}^\top\mathbf{c}_{1i})\left[\sum_{i=1}^{n_1}\boldsymbol{\mu}^\top\mathbf{D}_{1i}^\top\mathbf{V}_e^{-1}\mathbf{D}_{1i}\boldsymbol{\mu} + \sum_{i=1}^{n_1}\mathrm{Tr}(\mathbf{V}_e^{-1}\Sigma_i)\right]^{-1};$$

**until** *converged*;

**return** $(\boldsymbol{\alpha}/\lambda, \lambda^2\sigma_b^2, \sigma^2, \mathbf{V}_e)$ *after reduction step.*

### 2 TisCoMM-S$^2$

#### 2.1 TisCoMM model for summary statistics

To expand the applicability of TisCoMM to summary results of GWAS, in this section we extend our method for summary statistics (TisCoMM-S$^2$). We assume that the individual-level data $\mathcal{D}_2$ are not available, but instead the summary statistics from pair-wise marginal linear regression between $\mathbf{z}$ and $\mathbf{X}_2$ are provided:

$$\hat{\gamma}_j = (X_{2j}^\top X_{2j})^{-1} X_{2j}^T \mathbf{z}, \tag{S1}$$

$$\hat{\mathbf{s}}_j = (n X_{2j}^\top X_{2j})^{-1} (\mathbf{z} - X_{2j}\hat{\gamma}_j)^\top (\mathbf{z} - X_{2j}\hat{\gamma}_j), \tag{S2}$$

where $X_{2j}$ is the $j$th column of $\mathbf{X}_2$, $j \in \{1, \ldots, M\}$. In addition to the summary data, we also need the LD (correlations) matrix $R$ among those cis-SNPs. In practice, its estimate could be obtained from some database of genotypes in a suitable reference population. The computational details are discussed in the next section.

Let $\hat{\boldsymbol{\gamma}} = (\hat{\gamma}_1, \ldots, \hat{\gamma}_M)$, and $S = \text{diag}\{\mathbf{s}_1, \ldots, \mathbf{s}_M\}$. Following the similar argument of [5], it can be shown that

$$\hat{\boldsymbol{\gamma}} | S, R, \mathbf{b} \sim \mathcal{N}(SRS^{-1} \text{diag}\{\mathbf{b}\} \mathbf{W} \boldsymbol{\alpha}, \mathbf{V}), \tag{S3}$$

where $\mathbf{V} = SRS$.

To simplifies the computational requirements, we use a Cholesky decomposition to transform $\hat{\boldsymbol{\gamma}}$ to a vector of independent statistics. Obviously, $\mathbf{V}$ could be decomposed as

$$\mathbf{V} = \mathbf{L}^T \mathbf{L},$$

where $\mathbf{L}$ is an upper triangular $M \times M$ matrix with positive diagonal elements, and hence invertible. Transforming $\hat{\boldsymbol{\gamma}}$ by the Cholesky transpose inverse $(\mathbf{L}^\top)^{-1}$ we obtain a distribution of independent observations:

$$(\mathbf{L}^\top)^{-1}\hat{\boldsymbol{\gamma}}|S, R, \mathbf{b} \sim \mathcal{N}\left(\mathbf{U}\text{diag}\{\mathbf{b}\}\mathbf{W}\boldsymbol{\alpha}, \mathbf{I}_M\right)$$

where $\mathbf{U} = (\mathbf{L}^\top)^{-1}SRS^{-1}$.

Denote $\hat{\boldsymbol{\gamma}}^L = (\mathbf{L}^\top)^{-1}\hat{\boldsymbol{\gamma}}$. Now our model for GWAS summary statistic data become

$$\begin{aligned}
\mathbf{y}_i|\mathbf{b}, \mathbf{X}_1, \mathbf{W} &\sim \mathcal{N}\big(\sum_j x_{1ij} b_j \mathbf{w}_j, \mathbf{V}_e\big), \\
\hat{\boldsymbol{\gamma}}^L|\mathbf{U}, \mathbf{b} &\sim \mathcal{N}(\mathbf{U}\text{diag}\{\mathbf{b}\}\mathbf{W}\boldsymbol{\alpha}, \mathbf{I}_M), \\
\mathbf{b} &\sim \mathcal{N}(0, \sigma_b^2\mathbf{I}_M).
\end{aligned} \tag{S4}$$

Treating $(\mathbf{U}, \boldsymbol{\gamma}^L)$ as the pseudo individual data $\mathcal{D}_2$, the above model become to the individual data version of TisCoMM. Consequently, TisCoMM-S$^2$ can be inferred by an exactly similar procedure as the individual data version, except that there is no need to estimate $\sigma^2$ here. The TisCoMM joint test and tissue-specific test based on summary statistics also can be performed similarly.

### 3 Additional results for NFBC1966 study

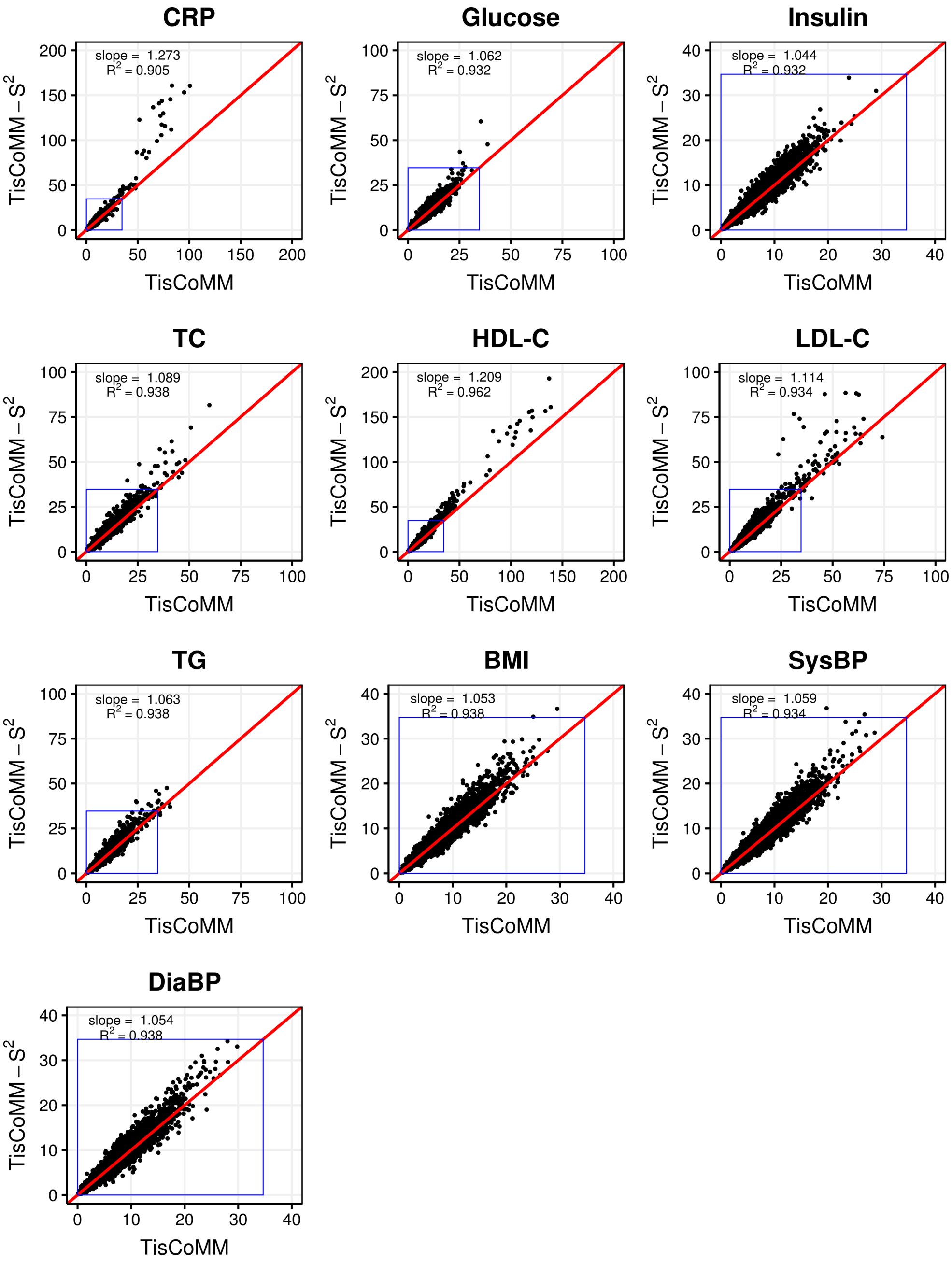

Figure S1: Comparison of TisCoMM and TisCoMM-$S^2$ results in NFBC1966 traits. The reference penal is European subsamples from 1000 Genomes Project. The blue rectangle indicates the null region. As the LD-structure between reference and study samples is mismatched, TisCoMM-$S^2$ becomes sightly inflated in the non-null regions.

### 4 Additional results for simulations

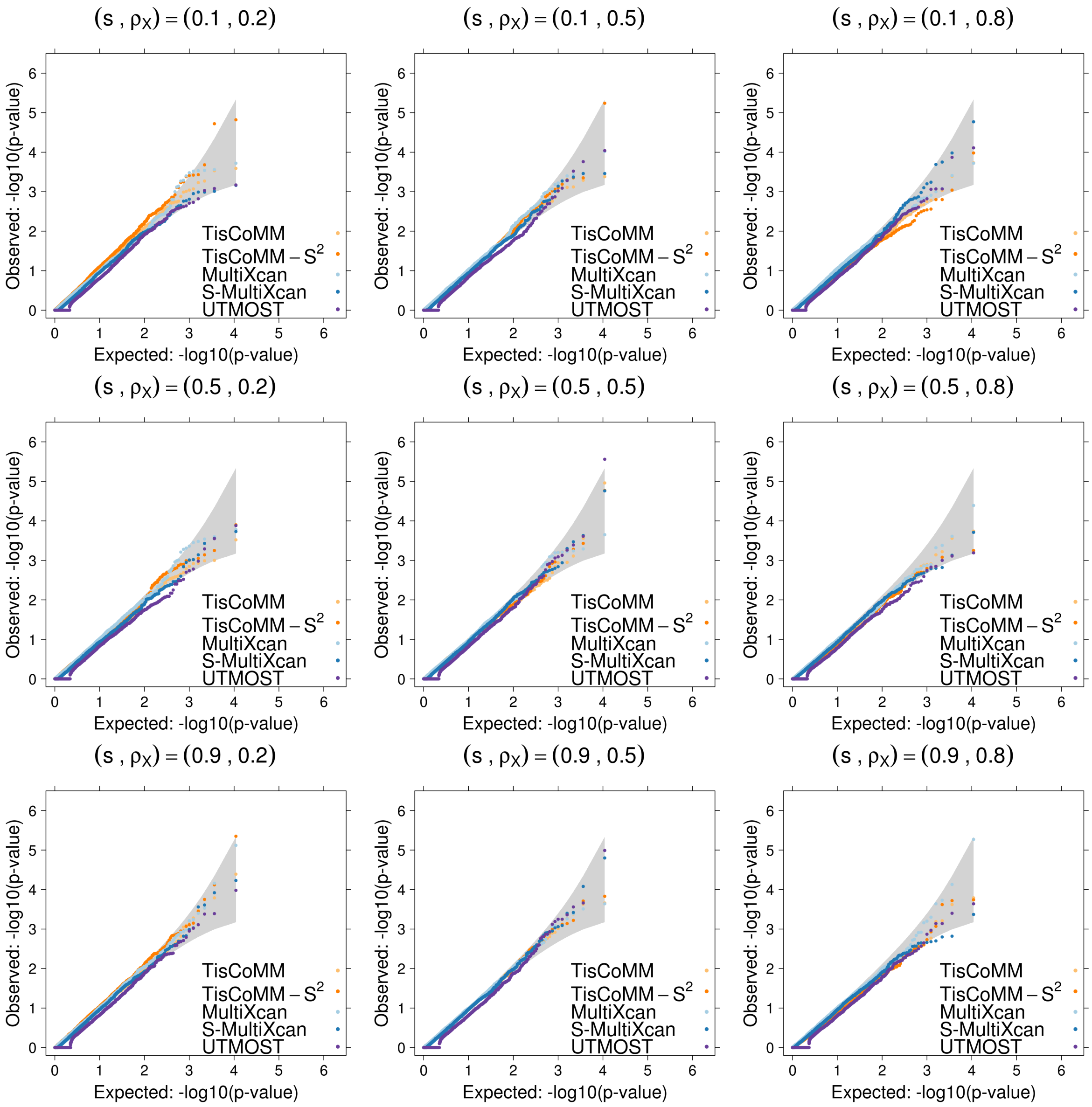

Figure S2: Quantile-quantile plot of $-log10$ p-values from different methods for testing gene-trait associations under the null models with different sparsity level $s$ and genotype correlation parameter $\rho_x$. The cellular heritability $h_c^2 = 0.025$.

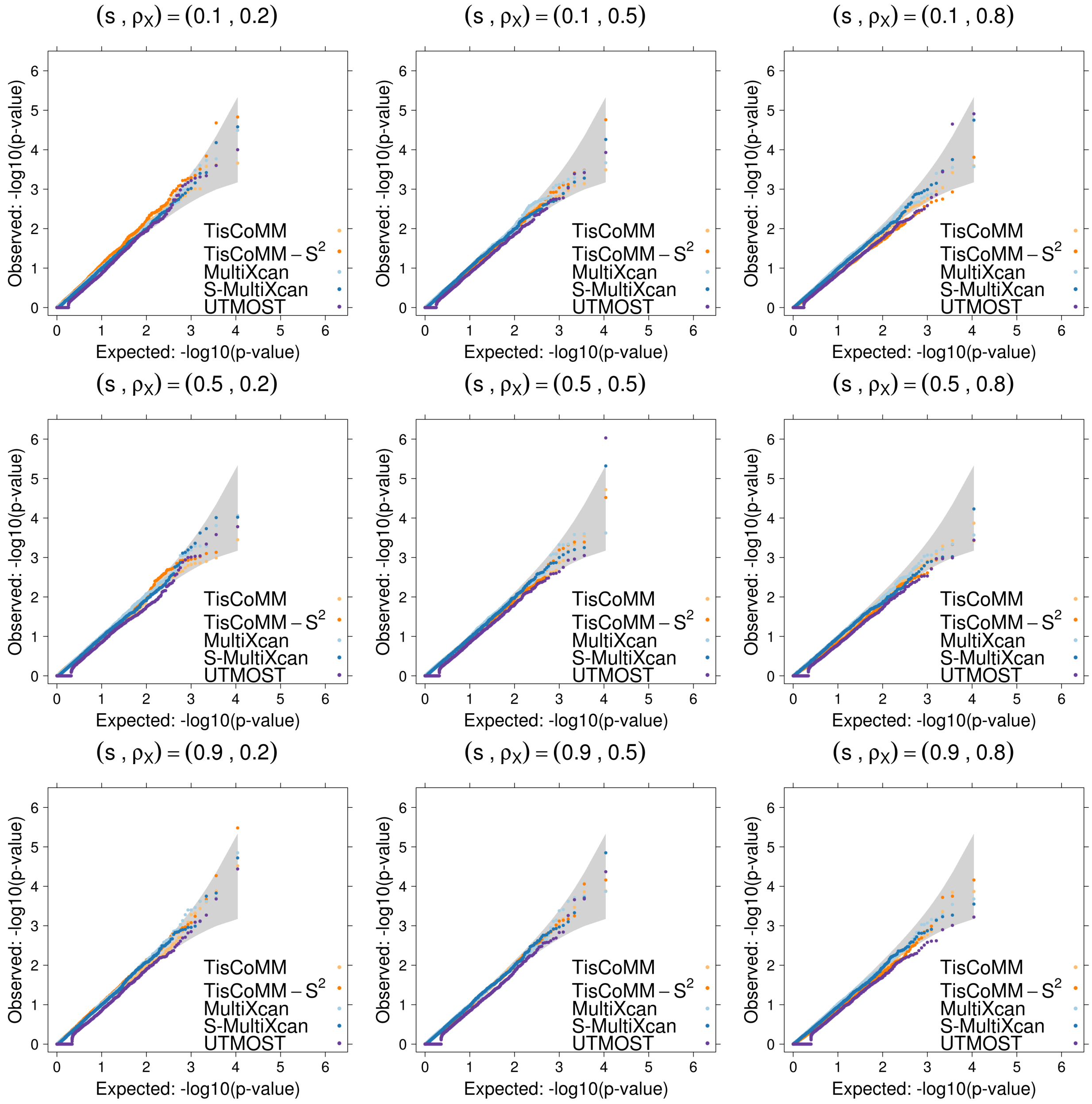

Figure S3: Quantile-quantile plot of $-log10$ p-values from different methods for testing gene-trait associations under the null models with different sparsity level $s$ and genotype correlation parameter $\rho_x$. The cellular heritability $h_c^2 = 0.05$.

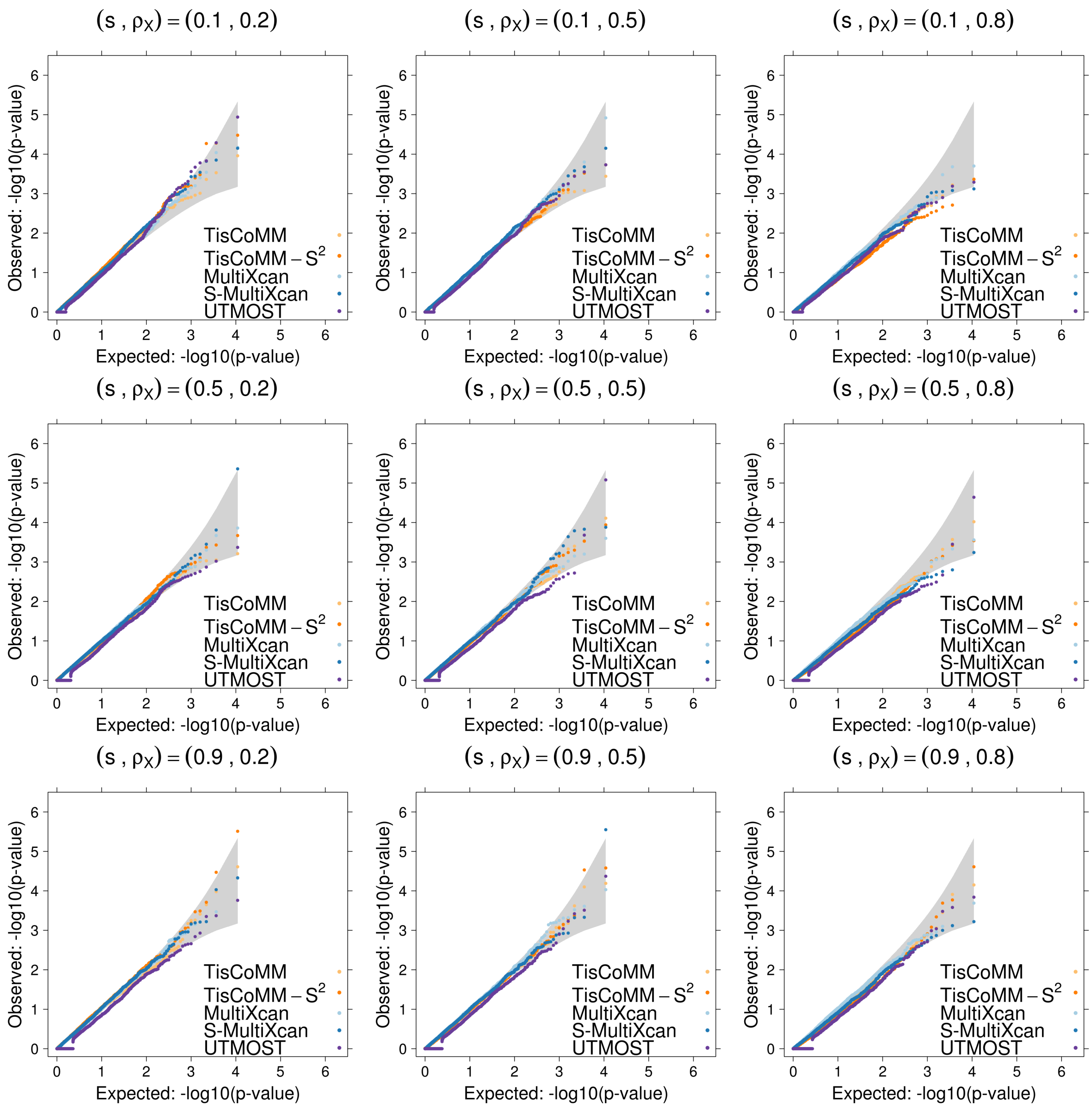

Figure S4: Quantile-quantile plot of $-log10$ p-values from different methods for testing gene-trait associations under the null models with different sparsity level $s$ and genotype correlation parameter $\rho_x$. The cellular heritability $h_c^2 = 0.1$.

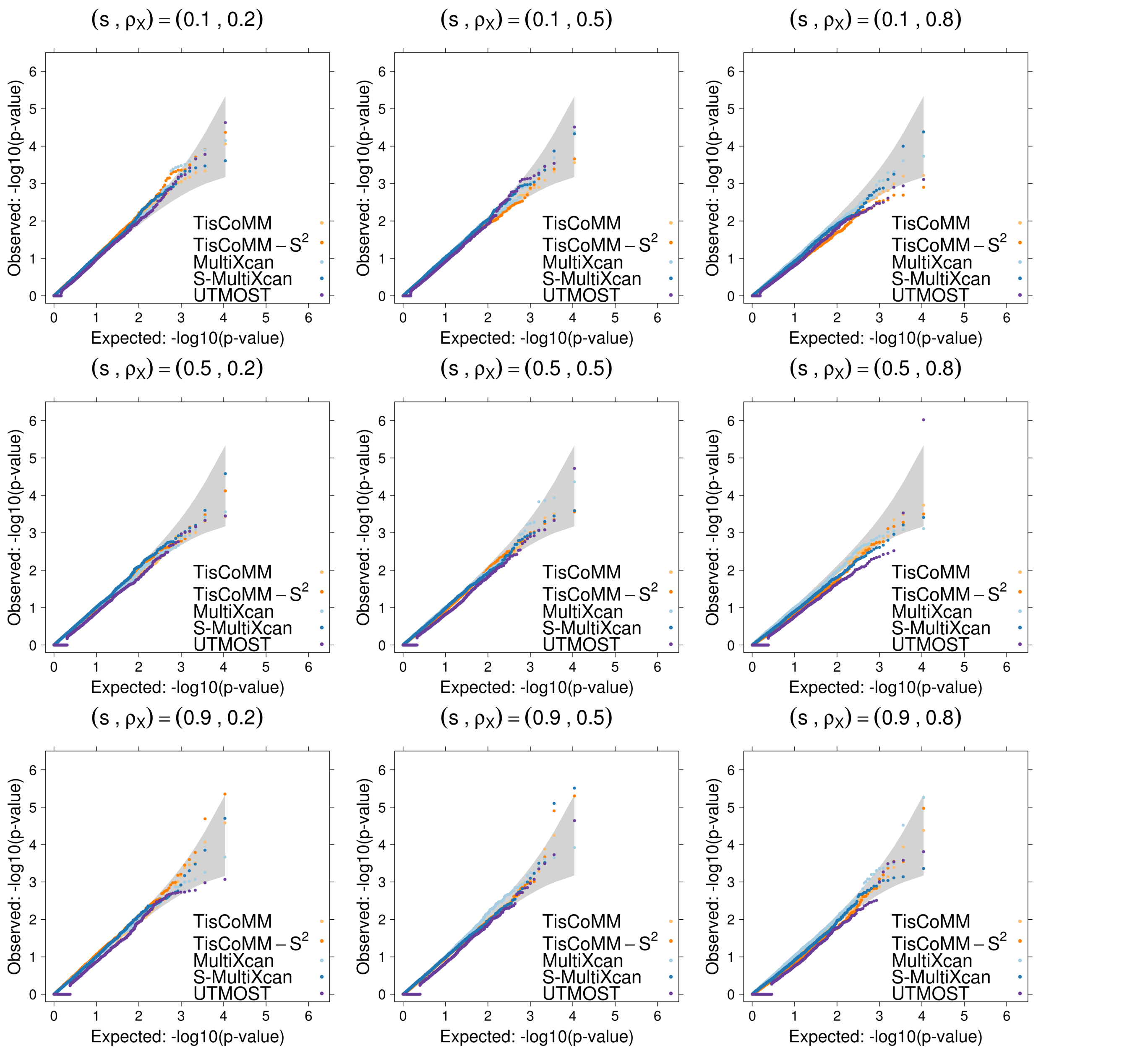

Figure S5: Quantile-quantile plot of $-log10$ p-values from different methods for testing gene-trait associations under the null models with different sparsity level $s$ and genotype correlation parameter $\rho_x$. The cellular heritability $h_c^2 = 0.2$.

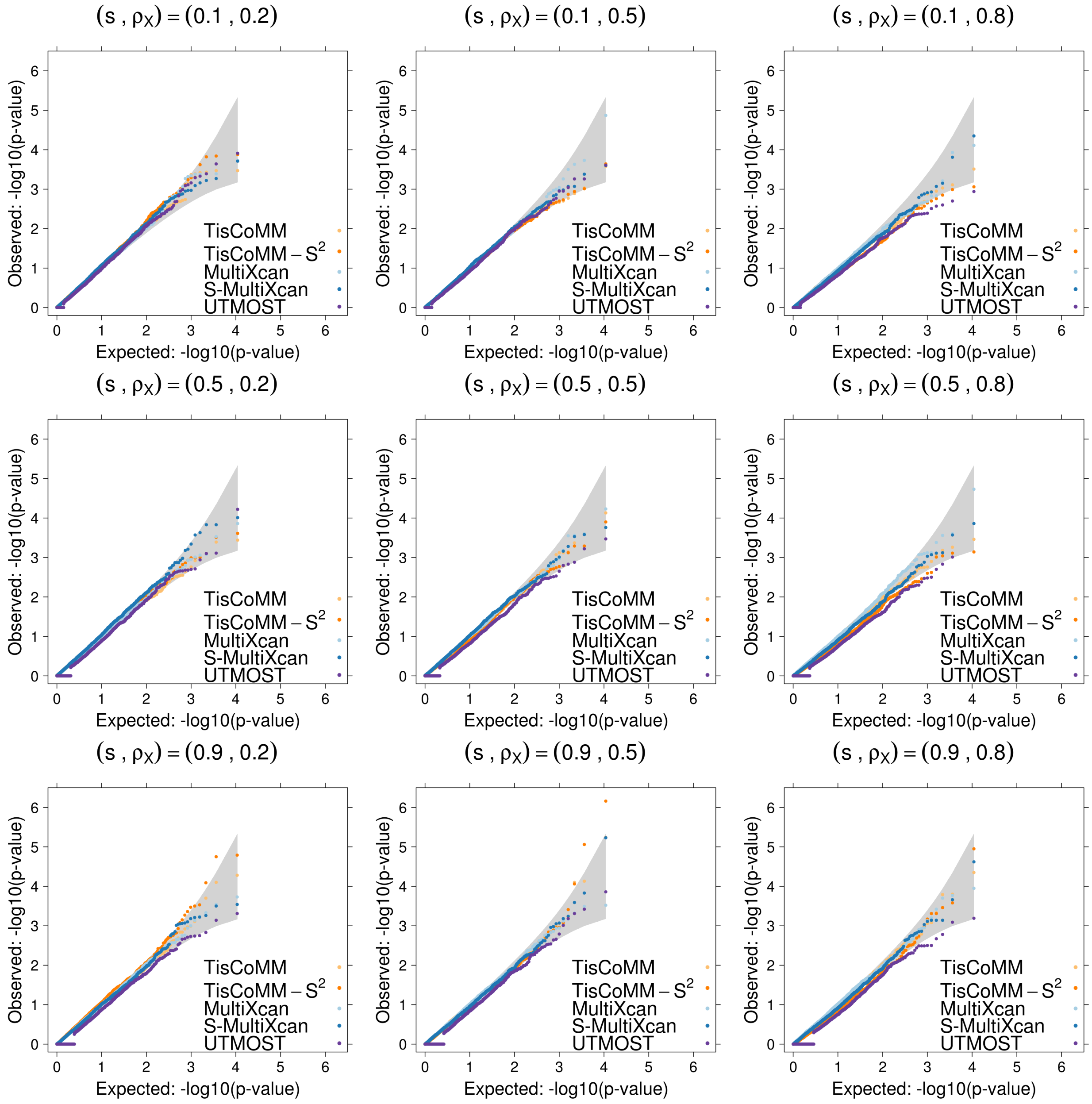

Figure S6: Quantile-quantile plot of $-log10$ p-values from different methods for testing gene-trait associations under the null models with different sparsity level $s$ and genotype correlation parameter $\rho_x$. The cellular heritability $h_c^2 = 0.4$.

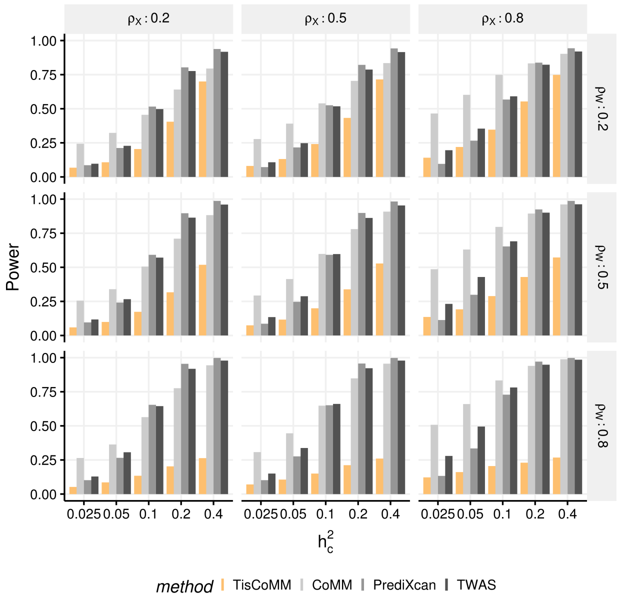

(A) Power

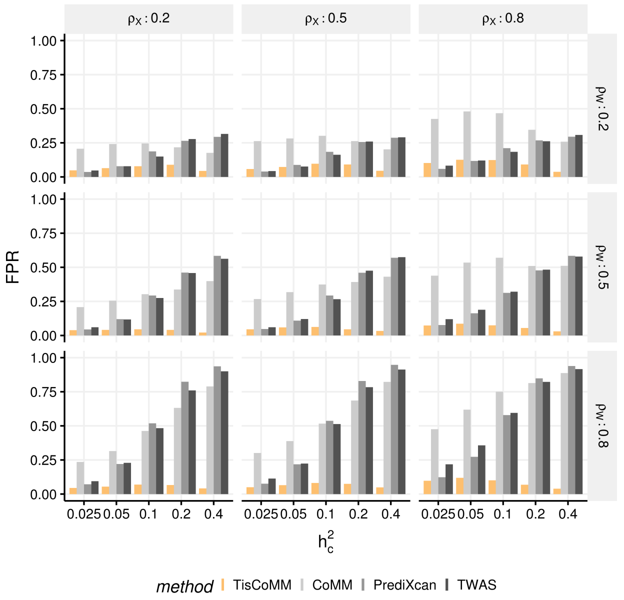

(B) false positive rate (FPR)

Figure S7: The comparison of TisCoMM tissue-specific test and the single-tissue association tests under the alternative hypothesis with two causal tissue. **A.** The power of TisCoMM tissue-specific test and the single tissue methods with Bonferroni correction applied. **B.** The corresponding false positive rates under each setting.

### 5 Additional results for NG and UKB traits

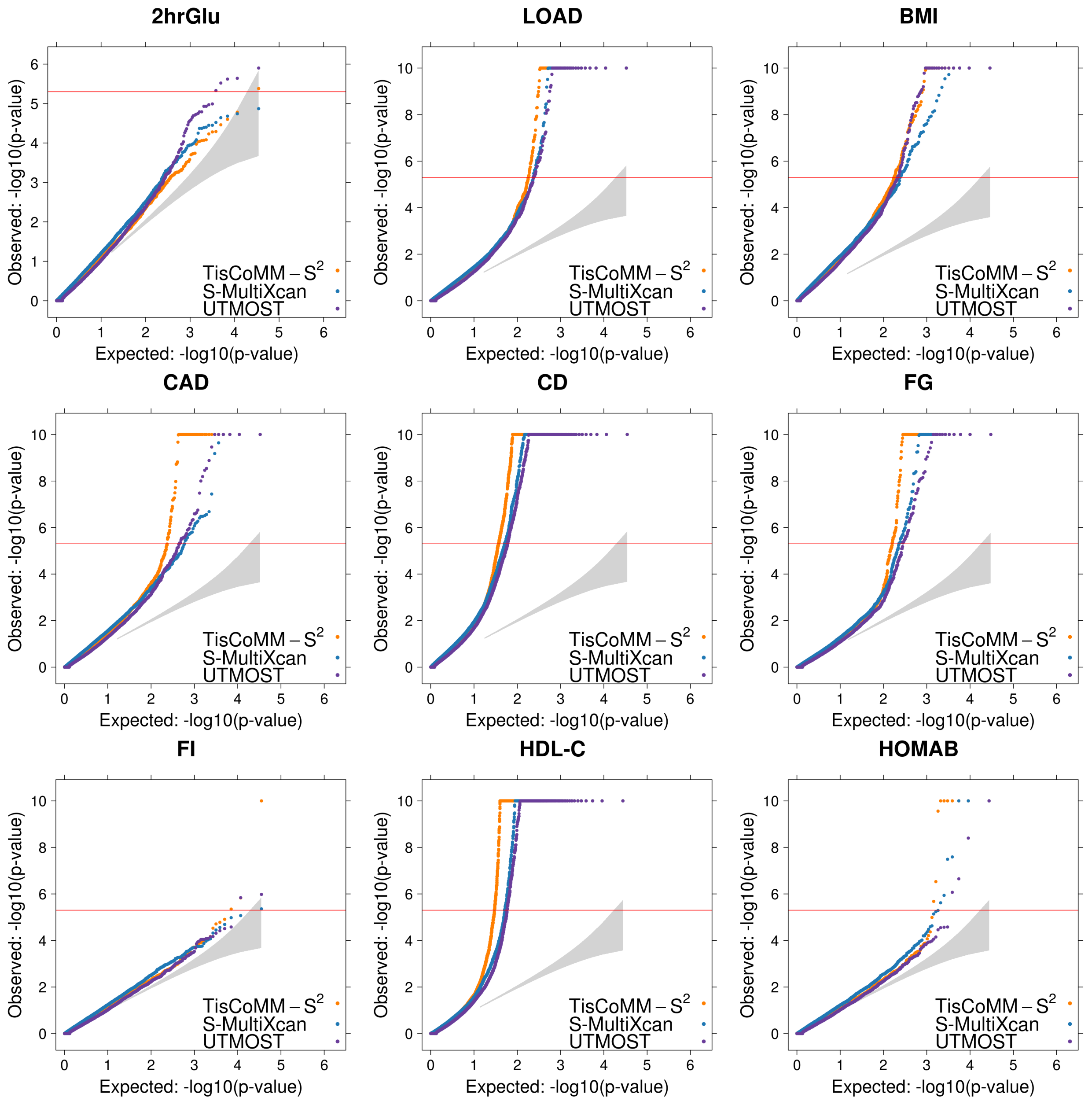

Figure S8: Quantile-quantile plots for 15 NG traits (Part I). The reference penal is European subsamples from 1000 Genomes Project.

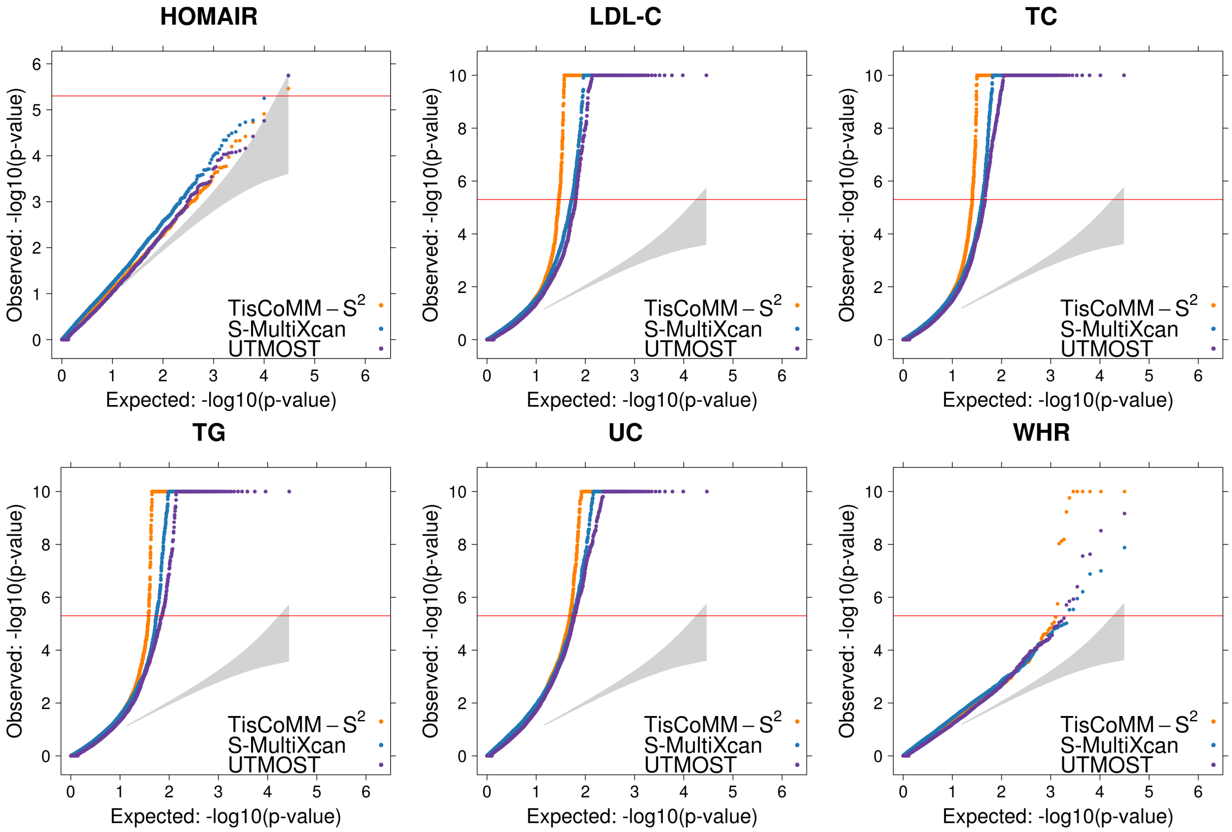

Figure S9: Quantile-quantile plots for 15 NG traits (Part II). The reference penal is European subsamples from 1000 Genomes Project.

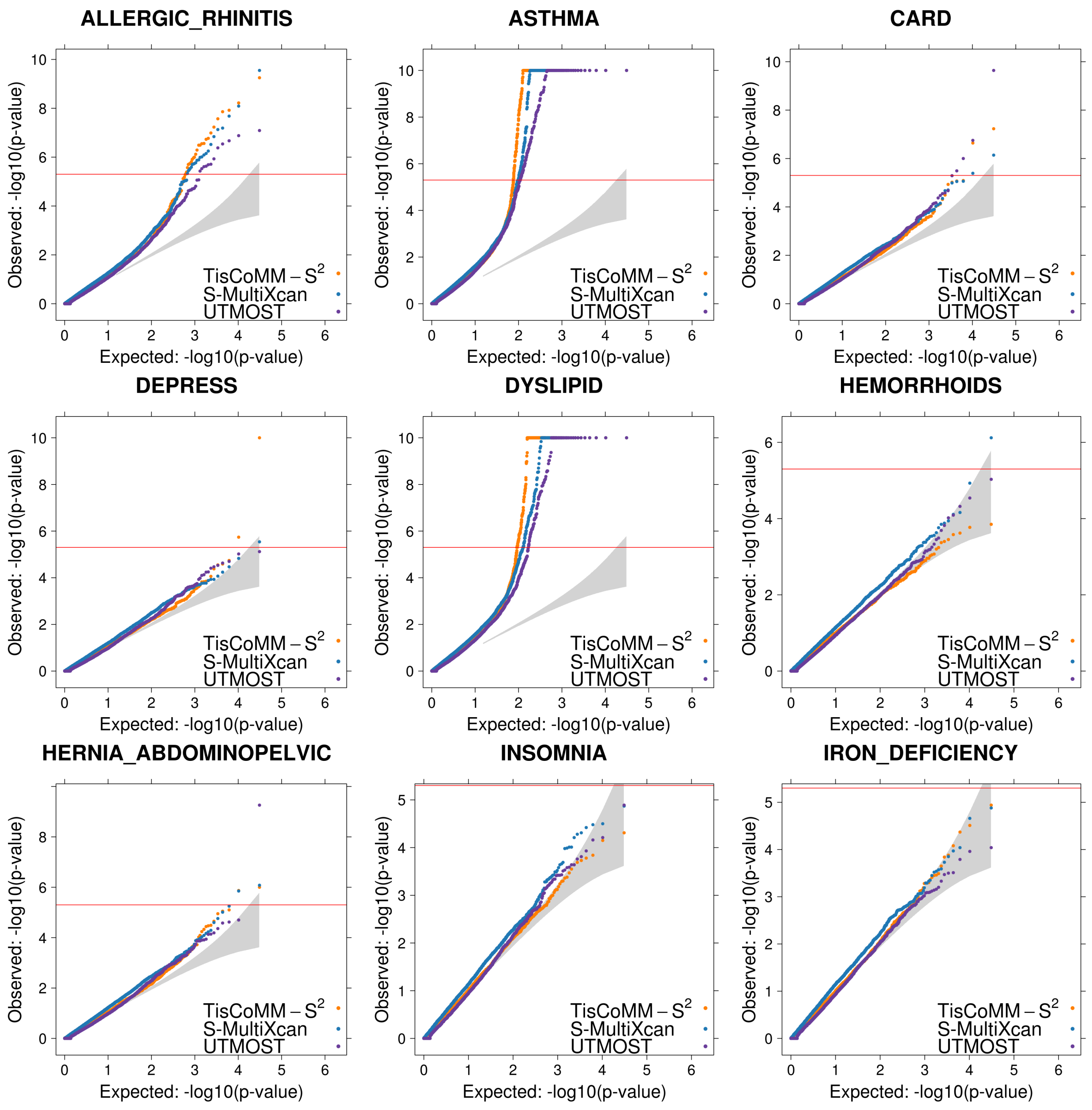

Figure S10: Quantile-quantile plots for 20 UKB traits (Part I). The reference penal is European subsamples from 1000 Genomes Project.

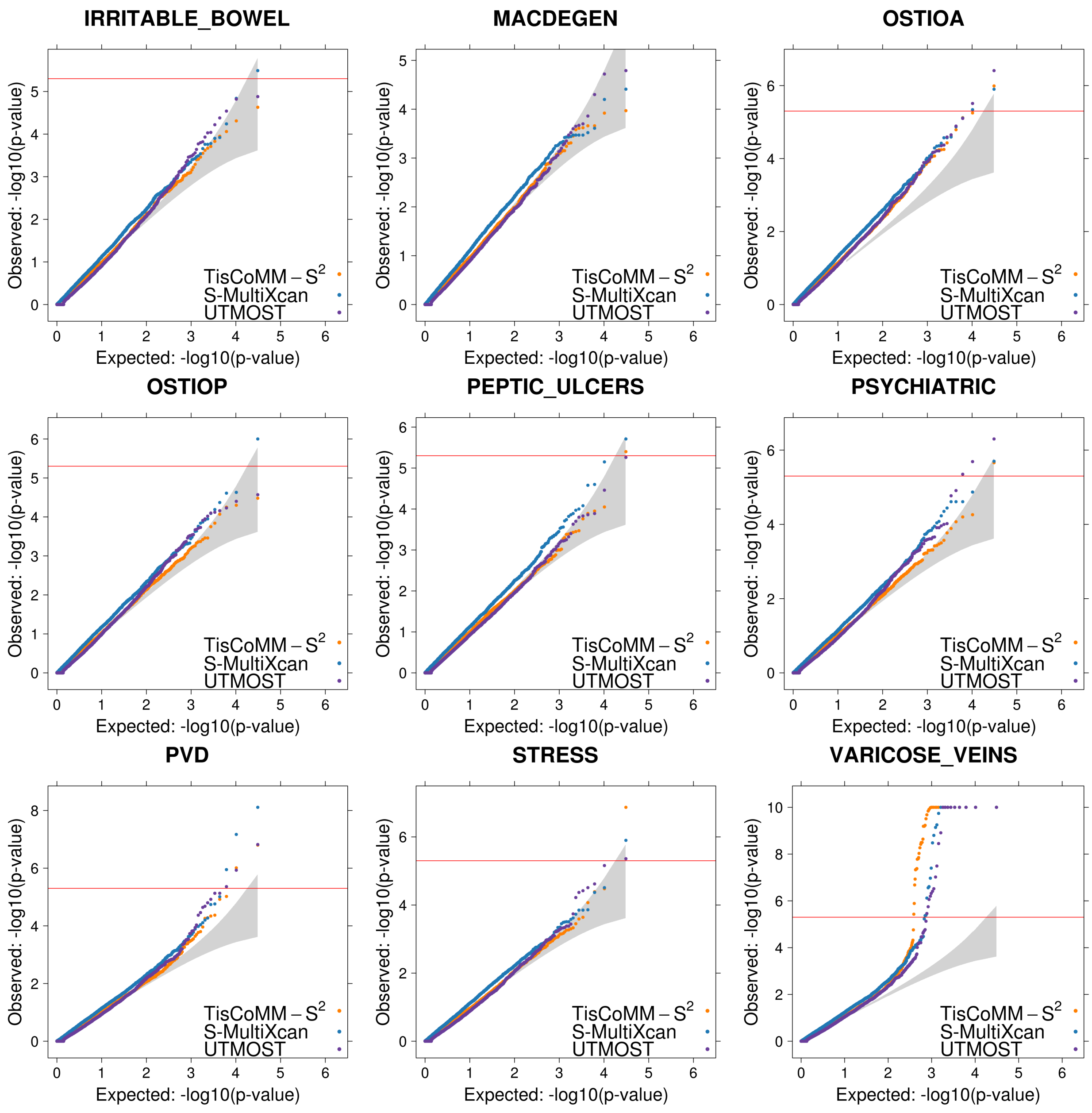

Figure S11: Quantile-quantile plots for 20 UKB traits (Part II). The reference penal is European subsamples from 1000 Genomes Project.

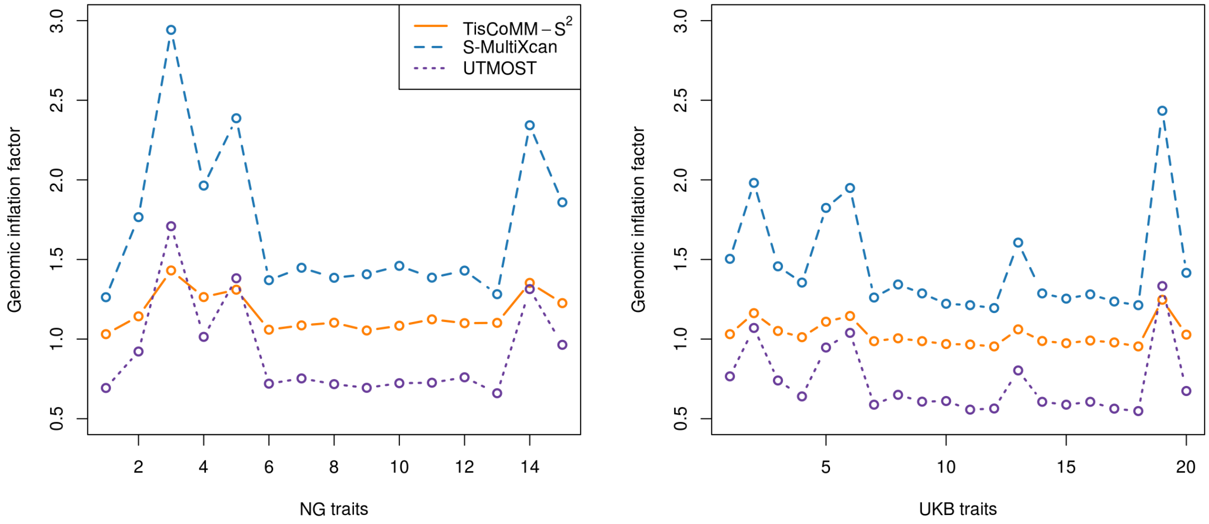

Figure S12: Genomic inflation factor for testing gene-trait associations for each of the NG and UKB traits by different methods. For TisCoMM-$S^2$, the genomic inflation factor is calculated from the the likelihood-ratio test statistics. For S-MultiXcan and UTMOST, their genomic inflation factors are calculated from p-values.

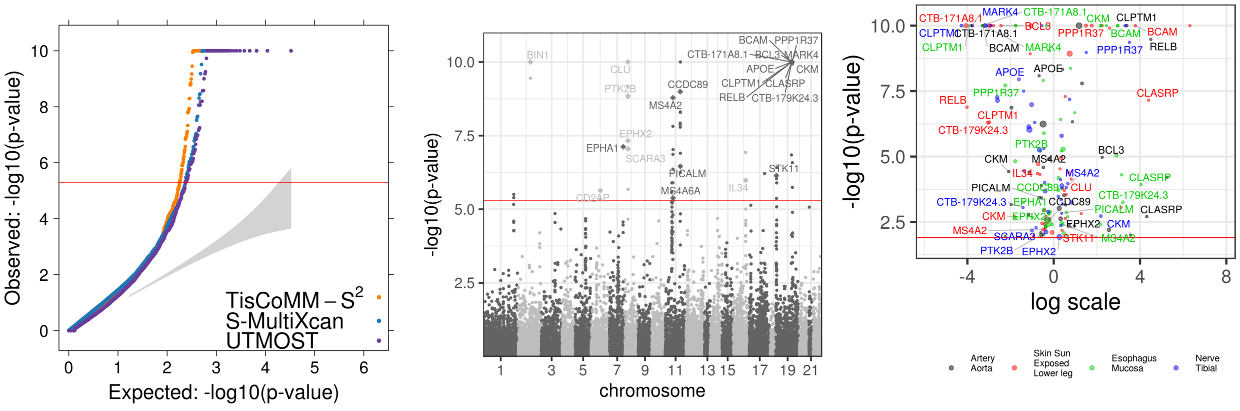

(A) LOAD

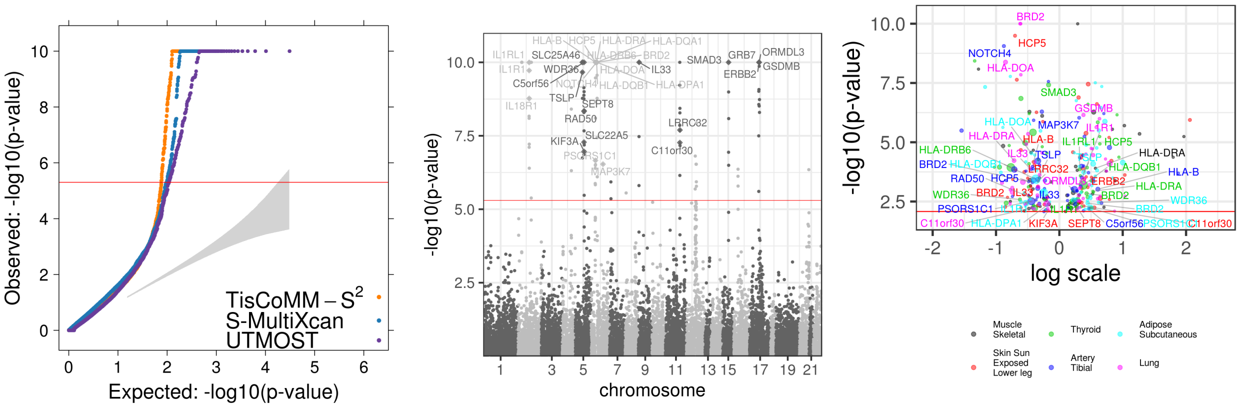

(B) Asthma

Figure S13: TisCoMM-S$^2$ results for LOAD and asthma. The reference panel is European subsamples from the 1000 Genomes Project. In each row, the three panels show the QQ plot (left), the Manhatton plot (middle), and the Volcano plot (right).

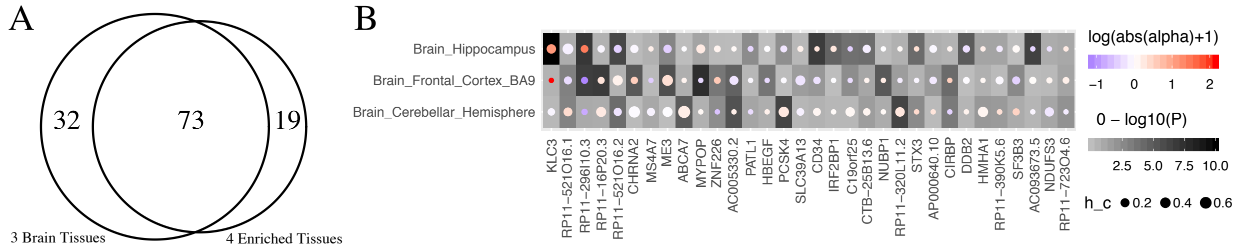

Figure S14: TisCoMM-S$^2$ results for LOAD based on three brain tissues. **A** Venn diagram showing the significant LOAD-associated genes shared by joint test based on three brain tissues and four enriched tissues. **B** Heatmap showing the results of the TisCoMM-S$^2$ tissue-specific test based on three brain tissues. The x-axis represents 32 LOAD-associated genes uniquely identified by the TisCoMM-S$^2$ joint test. The y-axis represents different tissue types. In each cell, the background color (shades of gray) indicates the significance level, the circle size indicates the heritability, and the color inside each circle indicates the effect size.

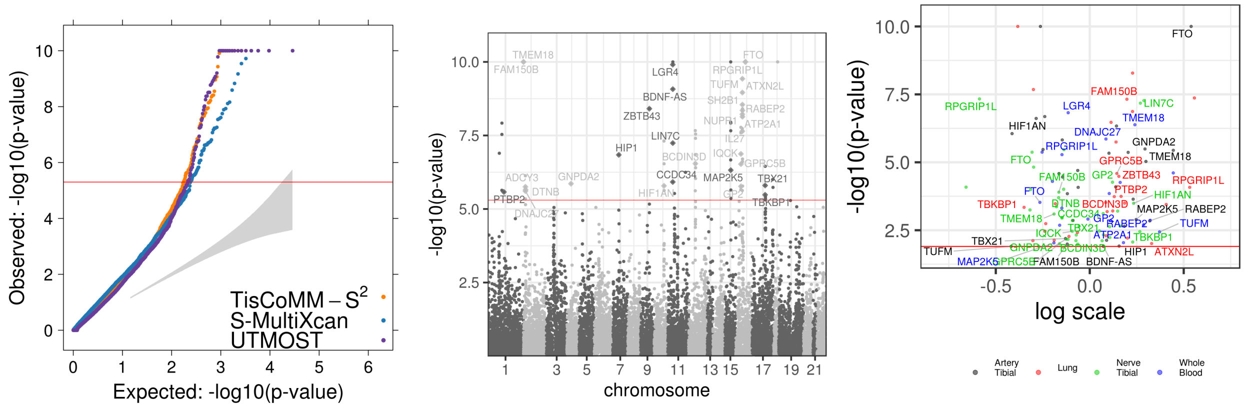

(A) BMI

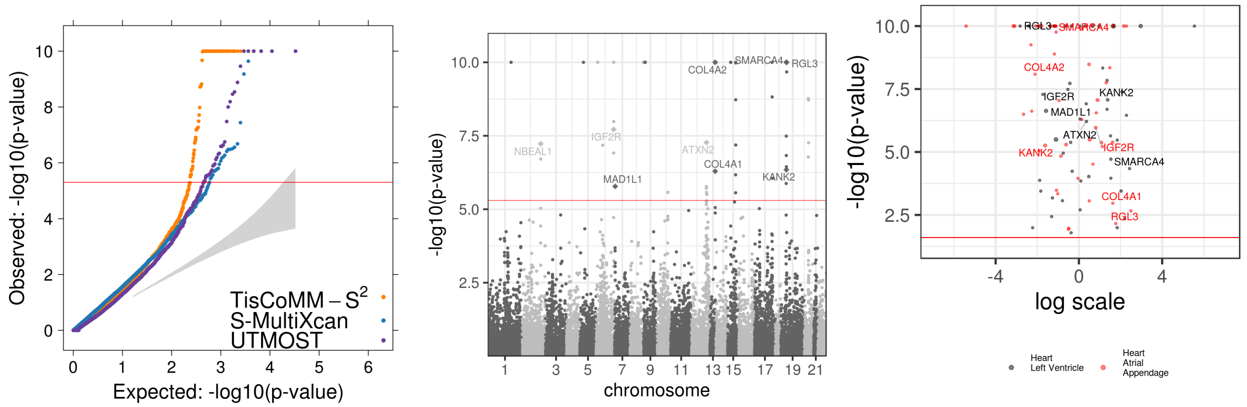

(B) CAD

Figure S15: TisCoMM-S$^2$ results for BMI and CAD. The reference panel is European subsamples from the 1000 Genomes Project. In each row, the three panels show the QQ plot (left), the Manhatton plot (middle), and the Volcano plot (right).

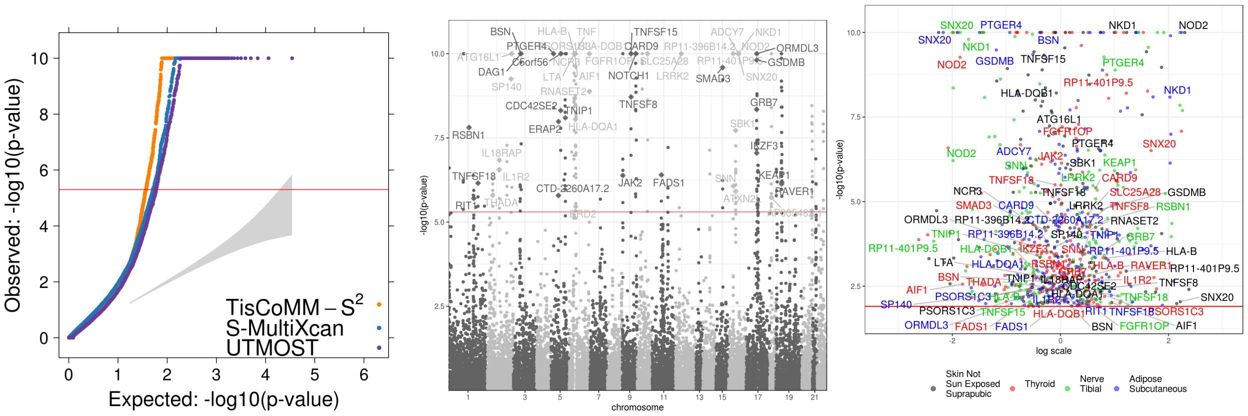

(A) CD

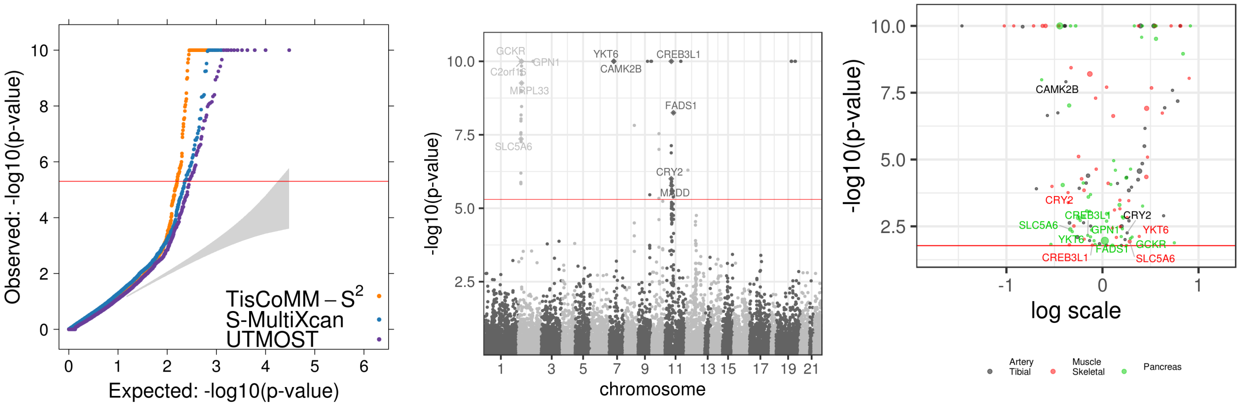

(B) FG

Figure S16: TisCoMM-S$^2$ results for CD and FG. The reference panel is European subsamples from the 1000 Genomes Project. In each row, the three panels show the QQ plot (left), the Manhatton plot (middle), and the Volcano plot (right).

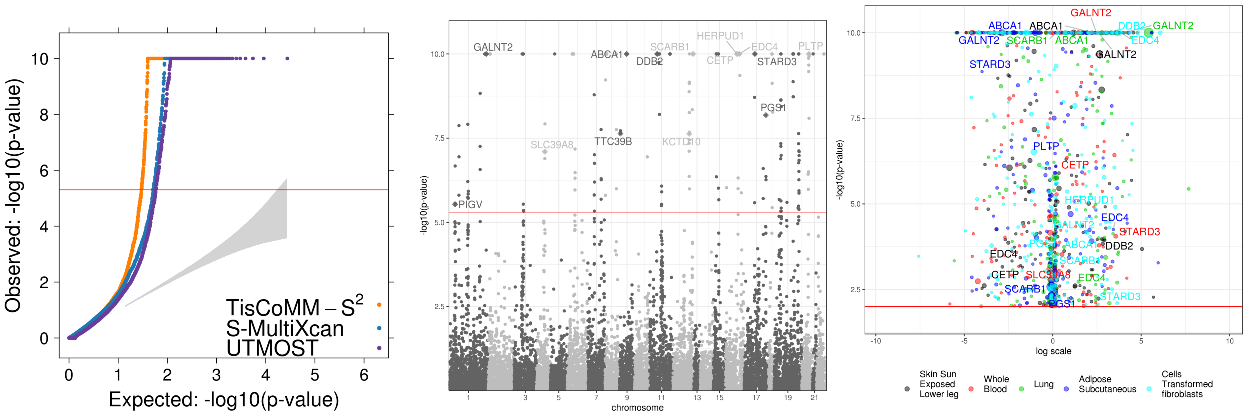

(A) HDL-C

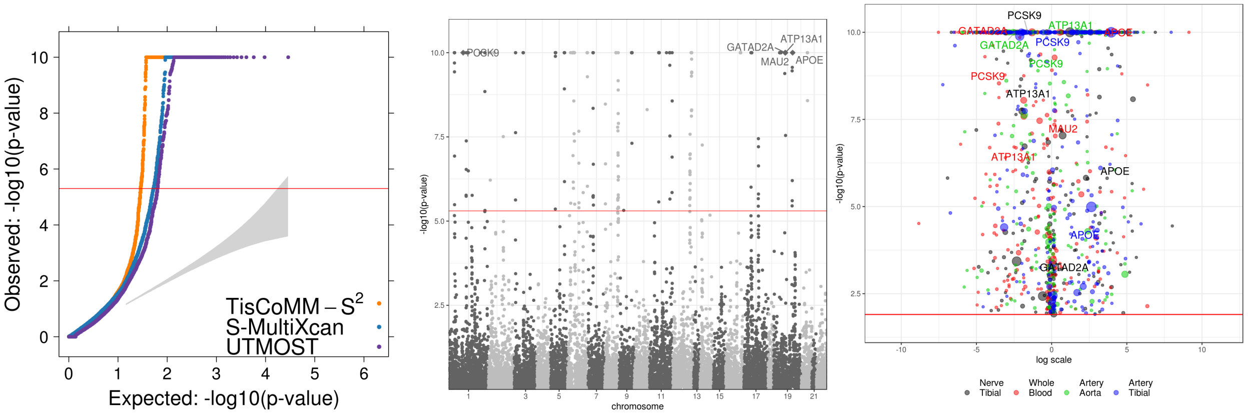

(B) LDL-C

Figure S17: TisCoMM-S$^2$ results for HDL-C and LDL-C. The reference panel is European subsamples from the 1000 Genomes Project. In each row, the three panels show the QQ plot (left), the Manhatton plot (middle), and the Volcano plot (right).

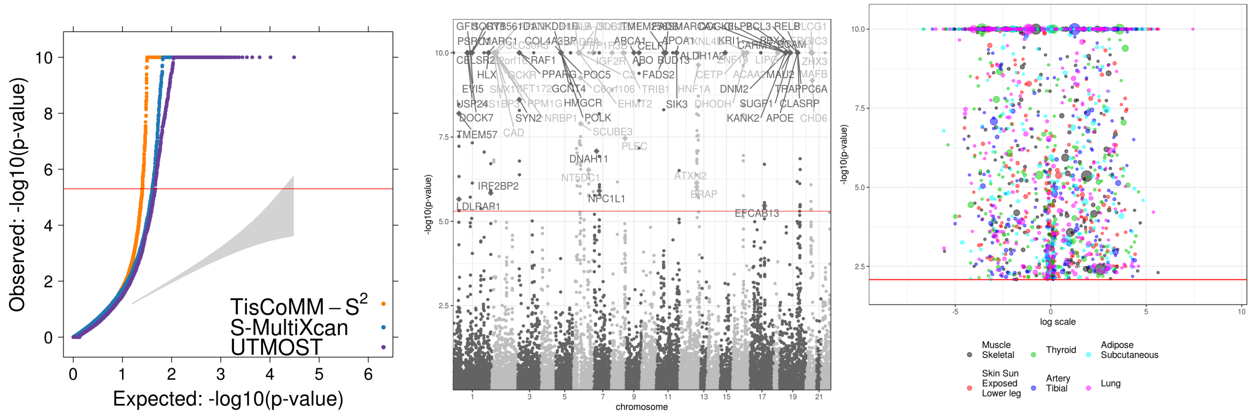

(A) TC

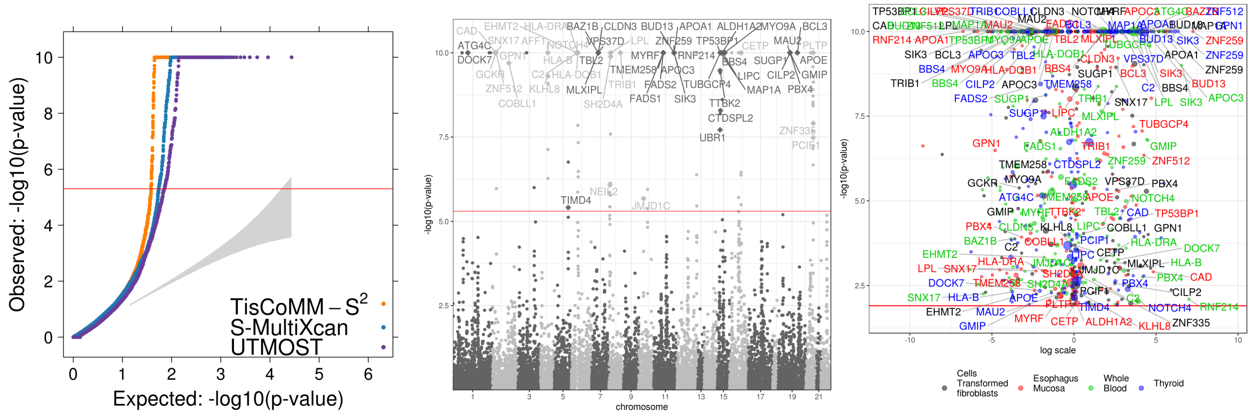

(B) TG

Figure S18: TisCoMM-S$^2$ results for TC and TG. The reference panel is European subsamples from the 1000 Genomes Project. In each row, the three panels show the QQ plot (left), the Manhatton plot (middle), and the Volcano plot (right). In the Volcano plot of TC, we do no annotate genes linked to TC because of the huge amount.

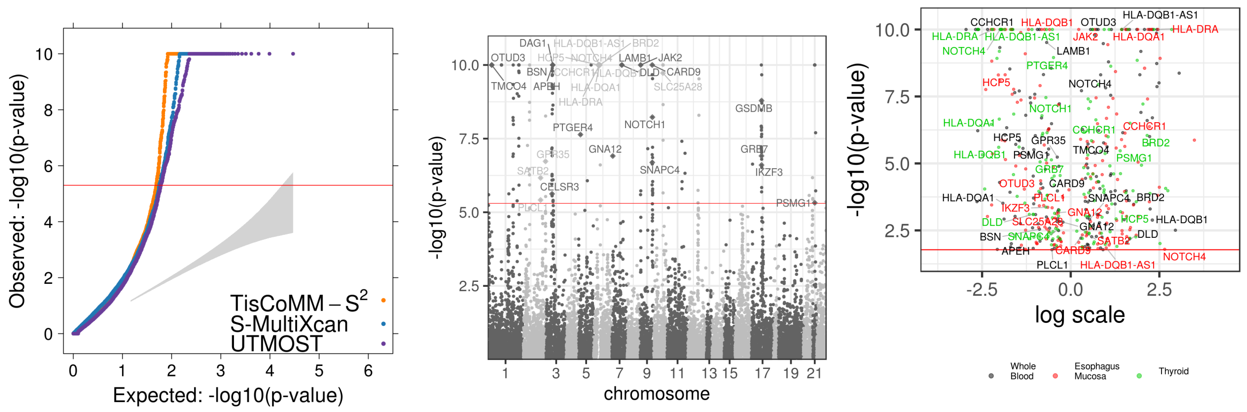

(A) UC

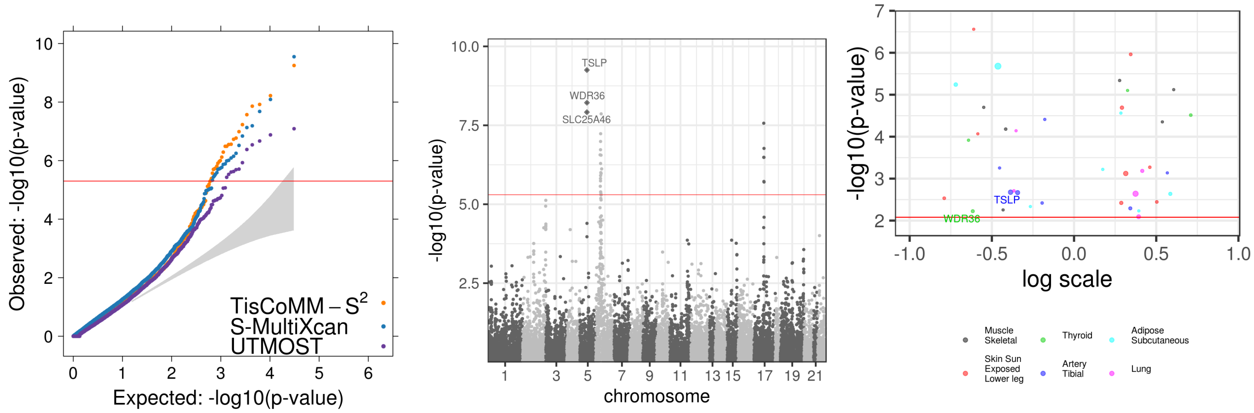

(B) ALLERGIC-RHINITIS

Figure S19: TisCoMM-$S^2$ results for UC and ALLERGIC-RHINITIS. The reference panel is European subsamples from the 1000 Genomes Project. In each row, the three panels show the QQ plot (left), the Manhatton plot (middle), and the Volcano plot (right).

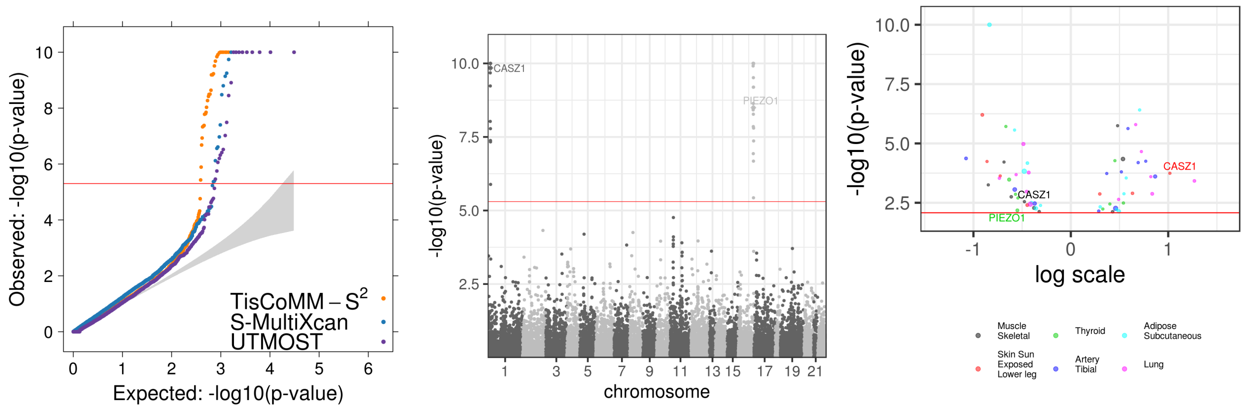

Figure S20: TisCoMM-S$^2$ results for DIA2 and VARICOSE-VEINS. The reference panel is European subsamples from the 1000 Genomes Project. In each row, the three panels show the QQ plot (left), the Manhatton plot (middle), and the Volcano plot (right).

5. Xiang Zhu and Matthew Stephens. Bayesian large-scale multiple regression with summary statistics from genome-wide association studies. *The annals of applied statistics*, 11(3):1561, 2017.
